# Contrasting and conserved roles of NPR pathways in diverged land plant lineages

**DOI:** 10.1101/2022.07.19.500630

**Authors:** Hyung-Woo Jeon, Hidekazu Iwakawa, Satoshi Naramoto, Cornelia Herrfurth, Nora Gutsche, Titus Schlüter, Junko Kyozuka, Shingo Miyauchi, Ivo Feussner, Sabine Zachgo, Hirofumi Nakagami

**Affiliations:** Max-Planck Institute for Plant Breeding Research, 50829 Cologne, Germany; Graduate School of Life Sciences, Tohoku University, Sendai 980-8577, Japan; Service Unit for Metabolomics and Lipidomics, Göttingen Center for Molecular Biosciences (GZMB), University of Göttingen, 37077 Göttingen, Germany; Division of Botany, Osnabrück University, 49076 Osnabrück, Germany; Department for Plant Biochemistry, Albrecht von Haller Institute for Plant Sciences and Göttingen Center for Molecular Biosciences (GZMB), University of Göttingen, 37077 Göttingen, Germany; School of Biological Science and Technology, College of Science and Engineering, Kanazawa University, Kakuma-machi, Kanazawa 920-1192, Japan; Department of Biological Sciences, Faculty of Science, Hokkaido University, Sapporo 060-0810, Japan

## Abstract

The NPR proteins function as salicylic acid (SA) receptors in *Arabidopsis thaliana*. AtNPR1 plays a central role in SA-induced transcriptional reprogramming whereby positively regulates SA-mediated defense. NPRs are found in the genomes of nearly all land plants. However, we know little about the molecular functions and physiological roles of NPRs in most plant species. Our phylogenetic and alignment analyses show that Brassicaceae NPR1-like proteins have characteristically gained or lost functional residues or motifs identified in AtNPRs, pointing to the possibility of a unique evolutionary trajectory for the Brassicaceae NPR1-like proteins that has resulted in peculiar functions. In line with this observation, we find that the only NPR in *Marchantia polymorpha*, MpNPR, is not the master regulator of SA-induced transcriptional reprogramming and negatively regulates bacterial resistance in this species. Interspecies complementation analysis indicated that the molecular properties of AtNPR1 and MpNPR are partially conserved, implying the diversification of NPR-associated pathways contributed to distinct roles of NPR in different species. The Mp*npr* transcriptome suggested potential roles of MpNPR in heat and far-red light responses. We identify both Mp*npr* and At*npr1-1* display enhanced thermomorphogenesis. NPRs and NPR-associated pathways clearly have evolved distinctively in diverged land plant lineages to cope with different terrestrial environments.

## Introduction

Land plants deal with various environmental fluctuations or stresses such as shadiness, heat, and drought. Likewise, land plants are constantly surrounded by various microbes and insects and have developed sophisticated immune systems to fight off a wide range of pathogenic organisms. Plant immune responses are controlled by diverse phytohormones such as salicylic acid (SA), jasmonic acid (JA), and ethylene (ET), among which SA has indispensable roles in both local and systemic protection against biotrophic and hemibiotrophic pathogens (Peng et al., 2021; Zhou and Zhang, 2020). In *Arabidopsis thaliana*, NONEXPRESSOR OF PATHOGENESIS-RELATED GENES1 (NPR1) is a central player in SA-mediated plant immunity. AtNPR1 is responsible for the majority of SA- or benzothiadiazole S-methyl ester (BTH; a functional analog of SA)-induced transcriptional reprogramming, positioning AtNPR1 as the master regulator of the regulatory defense network induced by SA in *A. thaliana* (Blanco et al., 2009; Wang et al., 2006). Consistent with this, the controlled expression of AtNPR1 in rice enhances resistance against the bacterial and fungal pathogens, *Xanthomonas oryzae* pv. *oryzae*, *Xanthomonas oryzae* pv. *oryzicola*, and *Magnaporthe oryzae*, without compromising plant fitness (Xu et al., 2017), underlining the potential of NPR homologs as molecular targets or tools for the engineering of disease-resistant plants.

AtNPR1 contains a Broad-complex, Tramtrack, and Bric-à-brac/poxvirus and zinc finger (BTB/POZ) domain; Ankyrin repeats (ANKs); and a C-terminal NPR1-like- C domain (Aravind and Koonin, 1999; Cao et al., 1997; Li et al., 2006; Ryals et al., 1997). The genome of *A. thaliana* encodes four NPR proteins, AtNPR1, AtNPR2, AtNPR3, and AtNPR4. Besides, *A. thaliana* has two BLADE-ON-PETIOLE (BOP) proteins, which are related to NPR proteins (Hepworth et al., 2005; Ha et al., 2004, 2007). Although AtNPR2 is closely related to AtNPR1, the At*npr2* mutant behaves like a wild type in response to SA, BTH, and bacterial pathogens (Castelló et al., 2018). However, expression of At*NPR2* can partially complement the SA- and BTH- induced resistance phenotype of the At*npr1-1* mutant (Castelló et al., 2018), suggesting functional conservation between AtNPR1 and AtNPR2 as positive regulators of SA signaling. Meanwhile, the At*npr3 npr4* double mutant displays enhanced resistance against bacterial and oomycete pathogens associated with elevated basal pathogen-related (*PR*) gene expression, and AtNPR3 and AtNPR4 have been shown to function as negative regulators of SA signaling (Ding et al., 2018; Zhang et al., 2006).

AtNPR1, AtNPR3, and AtNPR4 have been demonstrated to function as SA receptors. The SA-binding affinity of AtNPR proteins requires conserved arginine (Arg) residue in the C-terminal NPR1-like-C domains (Ding et al., 2018; Fu et al., 2012; Wu et al., 2012). An AtNPR1 mutant protein with the R432Q substitution failed to bind to SA, and the At*npr1-1* mutant expressing this protein could not complement the SA-insensitivity phenotype (Ding et al., 2018). Similarly, *Brachypodium distachyon* NPR2, BdNPR2, was shown to possess SA binding ability, which depends on the conserved Arg residue (Shimizu et al., 2022). AtNPR3 and AtNPR4 can function as transcriptional co-repressors, a function conferred by a repression motif, which is related to the ethylene-responsive element binding factor (ERF)- associated amphiphilic repression (EAR) motif (Ohta et al., 2001), found at the C- terminal NPR1-like-C domains of AtNPR3 and AtNPR4 but not AtNPR1 (Ding et al., 2018). Recent structural analysis suggested that SA binding to AtNPR1 leads to formation of an “enhanceosome” (Kumar et al., 2022). The structural basis of how AtNPR3 and AtNPR4 function as transcriptional co-repressors remains to be addressed.

AtNPR1 is a cysteine (Cys)-rich protein, and has been proposed to function as a redox sensor (Withers and Dong, 2016). In its inactive state, AtNPR1 forms oligomers through intermolecular disulfide bonds between Cys residues that act to sequester the protein in the cytoplasm (Mou et al., 2003). Importantly, Cys156 of AtNPR1 was shown to be targeted by S-nitrosylation, which facilitates oligomerization (Tada et al., 2008). Intriguingly, a recent study indicated that the disulfide-bridged oligomerization of AtNPR1, which was reported to occur *in vitro,* likely does not occur *in vivo* (Ishihama et al., 2021). Besides, AtNPR1, AtNPR3, and AtNPR4 were shown to regulate protein degradation by recruiting the Cullin 3 E3 ubiquitin ligase, likely through their BTB/POZ domains (Zavaliev et al., 2020).

The monophyletic bryophytes is a sister clade to tracheophytes (Leebens-Mack et al., 2019; Li et al., 2020; Puttick et al., 2018). The two plant lineages diverged from the common ancestor of the land plants, which evolved from streptophyte algae about 500 million years ago (Bowman et al., 2016; Strother and Taylor, 2018). Such taxonomic importance has placed bryophytes in a significant position in the research of plant evolution and terrestrialization. Bryophytes consist of three divisions: mosses, liverworts, and hornworts (Naramoto et al., 2022; Wickett et al., 2014). The genomes of model bryophytes, the moss *Physcomitrium patens*, the liverwort *Marchantia polymorpha*, and the hornwort *Anthoceros agrestis*, encode two, one, and no NPR homologs, respectively (Bowman et al., 2017; Lang et al., 2018; Li et al., 2020). Notably, expression of *P. patens* Pp*NPR1*/Pp*3c21_7570* can partially restore the *PR1* gene expression and disease resistance phenotypes of the At*npr1-1* mutant, suggesting that the molecular properties of NPR proteins in bryophytes and angiosperms are evolutionarily conserved at some level (Peng et al., 2017). Accumulating evidence shows that ectopic expression of At*NPR1* or NPR homologs in crops, including wheat, tomato, tobacco, and cotton, could alter disease resistance, further supporting functional conservation of NPR proteins in many plant species (Backer et al., 2019; Gao et al., 2013; Kumar et al., 2013; Lin et al., 2004; Makandar et al., 2006; Matthews et al., 2014; Parkhi et al., 2010b, 2010a; Potlakayala et al., 2007; Wally et al., 2009).

Most genetic studies on *NPR* genes that have investigated their physiological roles in different plant species have been carried out in *A. thaliana*. In one rare study in another plant species, in which the authors employed RNAi-based Os*NPR1* knockdown lines in rice, it was reported that Os*NPR1* is responsible for expression levels of 47% of BTH responsive genes (Sugano et al., 2010). However, it is not yet conclusive whether OsNPR1 functions as the master regulator of SA signaling as in *A. thaliana*, because the study was based on knockdown lines, and *O. sativa* has two other NPR proteins that may also contribute to SA signaling (Sugano et al., 2010). Therefore, it is still possible that genes other than NPR factors play central roles in SA signaling and that *NPR* genes primarily function in other pathways in plants. In this connection, AtNPR1 was shown to positively regulate cold acclimation in SA- and TGA-independent manners (Olate et al., 2018). Besides, AtNPR1 was described to suppress the unfolded protein response (UPR) independently from SA by interacting with the UPR regulators bZIP28 and bZIP60 (Lai et al., 2018). Here, we address the functional conservation of NPR proteins in plants by utilizing the model liverwort *M. polymorpha* that possesses a single NPR, MpNPR.

## Results

### Brassicaceae NPR1-like proteins possess a peculiar set of functionally important amino acid residues

Extensive studies of AtNPR proteins have identified protein motifs and amino acid residues associated with the molecular mechanisms of how AtNPR proteins induce or repress the expression of SA-responsive genes. To study functional conservation of NPR proteins across different plant species, we performed comprehensive sequence analyses of NPR and BOP homologs. Amino acid sequences of MpNPR (Mp1g02380) and AtBOP1 (At3g57130) were used for BLASTP search against 78 protein databases covering algae, bryophytes, lycophytes, monilophytes, gymnosperms, and angiosperms (Dataset S2). We identified 194 NPR and 123 BOP homologs in the genomes of 68 species covering the significant lineages of land plants (Dataset S1). No NPR and BOP homologs were found in streptophyte algae. We found that the exon-intron organization in the regions where the BTB/POZ domains are encoded is highly conserved between NPR and BOP families (Figure S2). The phylogenetic tree shows that NPR and BOP proteins from non-seed plants are closely related (Figure 1). This suggests that NPR and BOP families diverged in the early stage of terrestrialization and may share a common ancestral gene.

**Figure 1.**
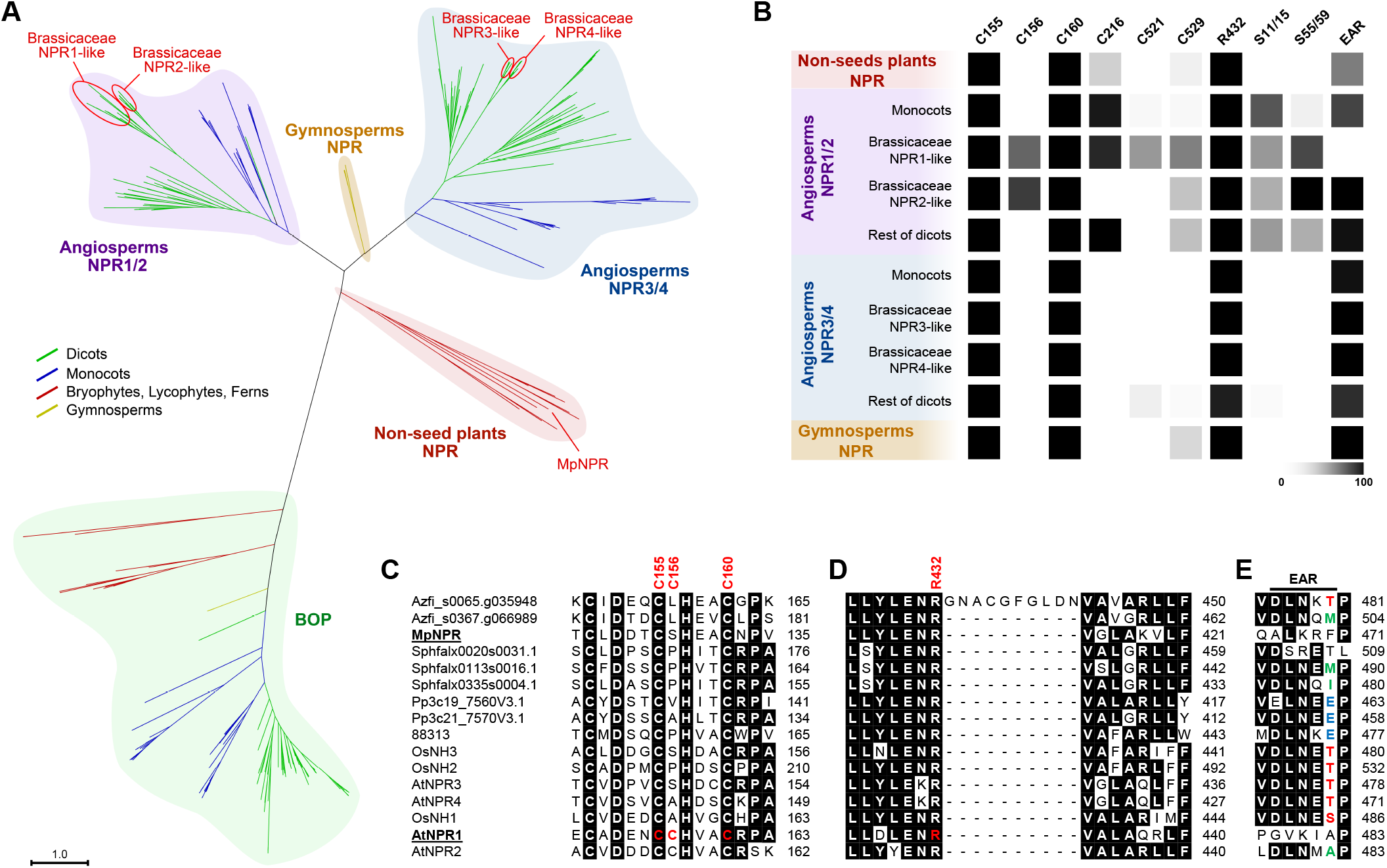
Phylogenetic analysis and amino acid conservation of 194 NPR and 123 BOP proteins from 68 plant species. (A) Unrooted phylogenetic tree showing that 194 NPR proteins were clustered into four major clades; angiosperms NPR1/2, angiosperms NPR3/4, non-seed plants NPR, and gymnosperms NPR clades. Brassicaceae species can be further classified into four subclades: Brassicaceae NPR1-like to Brassicaceae NPR4-like. (B–E) Conservation of functional amino acids and motifs associated with the molecular functions of AtNPR1. Cysteines required for AtNPR1 accumulation and oligomerization (C), Arg432 responsible for SA binding in AtNPR1 (D), and the EAR motif (DLNxxP) responsible for transcriptional co-repression activity (E) are shown.

Phylogenetic analysis of 194 NPR proteins classified the NPR family into three significant clades; Angiosperms NPR1/2, NPR3/4, and non-seed plants NPR (Figure 1A). The non-seed plants clade consists entirely of NPR homologs from bryophytes and pteridophytes. NPR homologs from angiosperms were clustered into the NPR1/2 and NPR3/4 clades, designated based on AtNPR1/AtNPR2 and AtNPR3/AtNPR4. The phylogenetic tree suggests that angiosperm NPR1/2 and NPR3/4 clades diverged from the ancestral NPR by gene duplication (Figure 1A). Our comprehensive analysis, including eight Brassicaceae species, revealed that *A. thaliana* gained a further four NPR proteins through additional gene duplication that occurred after the divergence of the Brassicaceae family (Figure 1A). Consequently, NPR proteins from Brassicaceae species can be further classified into four subclades: Brassicaceae NPR1-like to Brassicaceae NPR4-like (Figure 1A and S3B). Further expansions of *NPR1-like* genes were observed in *Boechera stricta*, *Brassica rapa FPsc*, and *Eutrema salsugineum*, while gene duplication of *NPR2-like*, *NPR3- like*, and *NPR4-like* was not observed except for *NPR3-like* in *Brassica rapa FPsc* (Figure S3A). Intriguingly, one of the *Boechera stricta* NPR1-like proteins, Bostr.29223s0069.1, was found to be integrated into the C-terminus of a Toll and IL-1 receptor (TIR)-NLR (Figure S3C), which implies that Brassicaceae NPR1-like proteins are virulence targets of pathogen effector molecules and, in turn, that, as supported by evidence from *A. thaliana*, Brassicaceae NPR1-like proteins play crucial roles in plant immunity.

We then investigated the conservation of amino acids such as cysteine (Cys), serine (Ser), and arginine (Arg) residues, which are associated with molecular functions of AtNPR1, together with functional motifs including the EAR motif and SUMO-interaction motif (SIM) (Figure 1B to 1E, and S1). AtNPR1 forms an oligomer through intermolecular disulfide bonds in its inactive state to reside in the cytoplasm, and several Cys residues were identified to be responsible for the oligomerization or protein accumulation (Mou et al., 2003). For instance, Cys156 is targeted by S-nitrosylation and is required for AtNPR1 oligomerization (Kumar et al., 2022; Tada et al., 2008). Cys521 and Cys529 are critical for transition metal binding activity, and oxidation is required for the SA-dependent co-activation activity of AtNPR1 (Rochon et al., 2006; Wu et al., 2012). We found that Cys155 and Cys160, required for accumulation of AtNPR1 protein (Mou et al., 2003), were conserved in all 194 NPR proteins (Figure 1B, 1C and S1). In contrast, Cys156 is conserved only in Brassicaceae NPR1-like and NPR2-like subclades (Figure 1B, 1C and S1). Cys82 is involved in the oligomerization of AtNPR1 (Mou et al., 2003; Kumar et al., 2022), and we found that Cys82 is well conserved among NPR proteins in all clades and could be detected in 185 out of 194 NPR proteins (Figure S1). Cys521 and Cys529 are modestly conserved only in Brassicaceae NPR1-like proteins (Figure 1B and S1). Arg432, required for SA binding in AtNPR1 (Ding et al., 2018), was found to be well conserved among NPR proteins in all clades and was detected in 190 out of 194 NPR proteins (Figure 1B, 1D and S1). The EAR motif (DLNxxP) of AtNPR3 and AtNPR4 was found to be responsible for their transcriptional co-repressor activities (Ding et al., 2018). Conversely, AtNPR1 was assumed to function as a transcriptional co-activator due to the absence of this motif (Innes, 2018). To our surprise, we found that the EAR motif is conserved in most NPR proteins except for the Brassicaceae NPR1-like proteins (Figure 1B, 1E and S1). Notably, the EAR motif is well conserved in Brassicaceae NPR2-like, NPR3-like, and NPR4-like proteins (Figure 1B and S1). It is likely that Brassicaceae NPR1-like proteins have lost the EAR motif during evolution. We observed occasional independent losses of the EAR motif in NPR proteins from other plant species, including the single NPR in *M. polymorpha*, MpNPR (Figure 1E and S1). In short, Brassicaceae species have a unique set of NPR proteins, and Brassicaceae NPR1-like proteins may have evolved unique molecular functions as positive regulators of SA-mediated immunity.

### NPR negatively regulates the SA response and resistance against bacterial pathogens in M. polymorpha

Based on the phylogenetic relationships of NPR proteins, functional characterization of NPR proteins from bryophytes would be expected to shed light on the ancestral function of NPR and its evolution. The acquisition of NPR has probably contributed to plant terrestrialization. However, the absence of *NPR* genes in those hornworts for which genomes have been sequenced, namely, *Anthoceros agrestis*, *Anthoceros punctatus*, and *Anthoceros angustus*, indicates that NPR is not indispensable for the survival of plants in terrestrial environments. The model liverwort *M. polymorpha* has a single NPR, which is an advantage for efforts aimed at addressing NPR functions. Since AtNPR1 plays a prominent and vital role in SA responses in *A. thaliana*, a comparison of MpNPR and AtNPR1, both of which lack the EAR motif, can contribute to understanding the conservation and diversification of NPR functions. We thus generated CRISPR/Cas9-based NPR loss-of-function mutants in the *M. polymorpha* Tak-1 background. Two guide RNAs were used to target the first exon of Mp*NPR* near the region encoding the BTB/POZ domain and resulted in 26-bp deletion (Mp*npr-1^ge^*) and 1-bp insertion (Mp*npr-2^ge^*), respectively (Figure S4A). Both alleles resulted in a frameshift and early stop codons. We did not observe obvious developmental defects of the Mp*npr* mutants under our growth conditions (Figure 2A).

**Figure 2.**
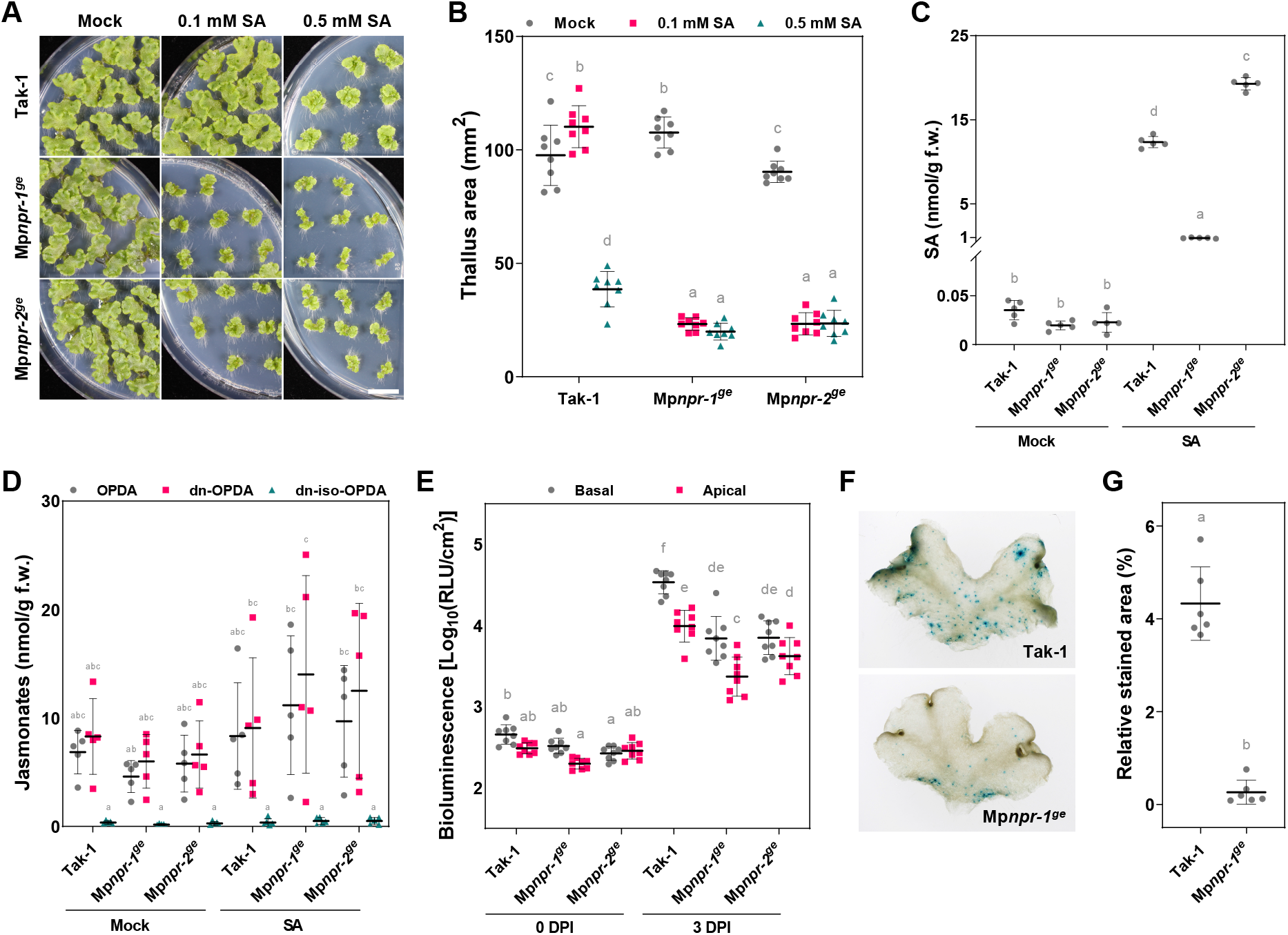
Phenotypic analyses of Mp*npr* mutants in SA and SA-induced immune responses. (A, B) SA hypersensitivity in the growth of Mp*npr-1^ge^* and Mp*npr-2^ge^* compared to Tak-1. Scale bar, 1 cm (n = 8). (C, D) SA and jasmonate measurements in Tak-1 and Mp*npr* mutants grown on mock and SA-supplemented conditions (n = 5). (E) Quantification of bacterial growth in the basal and apical thallus, inoculated with the bioluminescent *Pto*-lux (n = 8). DPI, days post-inoculation. (F–G) Transient transformation of Tak-1 and Mp*npr-1^ge^* using *Agrobacterium* carrying *intron-GUSPlus* (n = 6). (B–E, G) Different letters represent statistical significances (Tukey’s test; *p* < 0.01). Error bars indicate SD.

SA was shown to inhibit the growth of *M. polymorpha*, and thus we first investigated SA sensitivity of the Mp*npr* mutants (Gimenez-Ibanez et al., 2019). Tak- 1 plants showed no responses in their growth at concentrations up to 0.5 mM SA, while both Mp*npr* mutants displayed growth inhibition only at 0.1 mM SA (Figure 2A and 2B). Complemented alleles expressing Mp*NPR* under its own promoter in Mp*npr-1^ge^* partially rescued the SA hypersensitivity phenotype (Figure 5A, 5B, S4B and S4C). Moreover, Mp*NPR*-overexpressing plants, *_pro_*Mp*EF1*α:Mp*NPR- mCitrine/*Tak-1, displayed reduced sensitivity to the SA treatment (Figure 5A and 5B). At*npr1* mutants are characterized by an SA-hypersensitive phenotype due to hyperaccumulation of SA in these mutants (Zhang et al., 2010a). Therefore, we next measured SA levels in the Mp*npr* mutants to investigate whether the differential SA accumulation contributed to the hypersensitive phenotype. Under our normal growth conditions, SA accumulated to the same levels in the Tak1 and Mp*npr* mutants (Figure 2C). When plants were grown under SA-supplemented conditions, we observed significantly lower and higher SA levels in Mp*npr-1^ge^* and Mp*npr-2^ge^* compared to Tak-1, respectively. However, there was no correlation between the altered SA levels and the growth inhibition phenotype (Figure 2A to 2C). In *M. polymorpha*, *dinor*-12-oxo-phytodienoic acid (dn-OPDA) functions as bioactive jasmonate (JA), and antagonism between SA and dn-OPDA was reported (Gimenez-Ibanez et al., 2019; Matsui et al., 2020; Monte et al., 2018). Therefore, we also measured OPDA levels and found that these were not altered in the Mp*npr* mutants (Figure 2D). Altogether, we concluded that MpNPR negatively regulates SA-dependent growth inhibition.

**Figure 5.**
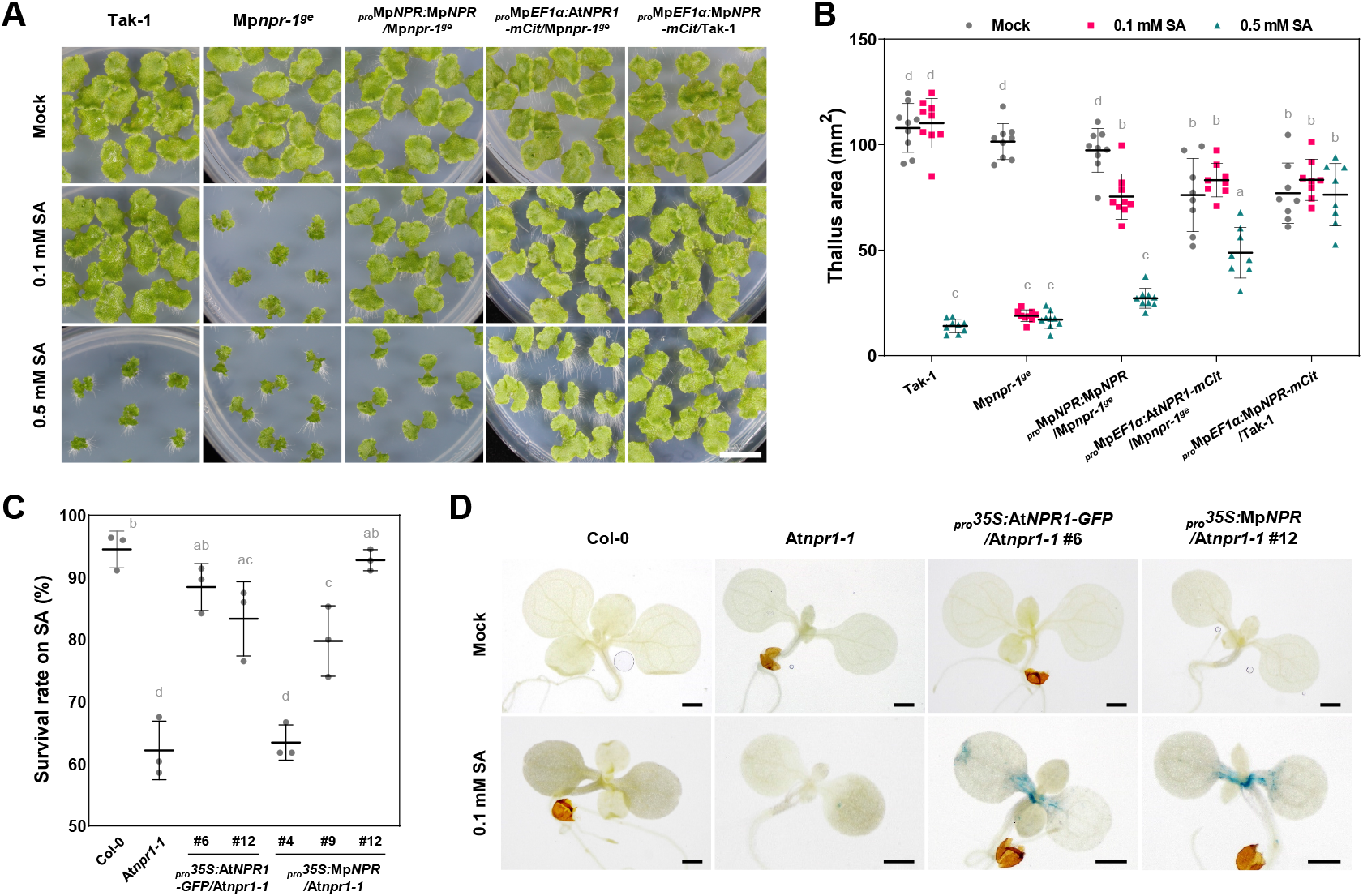
Inter-species complementation analysis in *M. polymorpha* and *A. thaliana*. (A, B) Complementation of SA hypersensitivity in Mp*npr-1^ge^* by introducing *_pro_*Mp*NPR:*Mp*NPR* and *_pro_*Mp*EF1α:*At*NPR1-mCitrine*. Scale bar, 1 cm (n = 8). (C) Complementation of SA hypersensitivity in At*npr1-1* by expression of *_pro_35S:*Mp*NPR* (n = 3). (D) GUS staining results showed that Mp*NPR* overexpression restored SA-induced *BGL2::GUS* expression in At*npr1-1*. Scale bars, 1 mm. (B, C) Different letters represent statistically significant differences (Tukey’s test; *p* < 0.01). Error bars indicate SD.

AtNPR1 positively regulates SA-dependent defense gene expression, and thus At*npr1* mutants display increased susceptibility to various bacterial pathogens (Backer et al., 2019; Cao et al., 1997; Roetschi et al., 2001). Therefore, we investigated whether and how MpNPR contributes to resistance against bacterial pathogens using recently established pathosystems (Iwakawa et al., 2021; Matsumoto et al., 2022). Two-week-old thalli of Tak-1 and Mp*npr* mutants were inoculated with bioluminescent *Pseudomonas syringae* pv. *tomato* DC3000 (*Pto*-lux), and bacterial growth was measured at 3 days post-inoculation (DPI). *Pto*-lux growth in the Mp*npr* mutants was significantly reduced compared to the growth in Tak-1 in both basal and apical regions of thalli (Figure 2E). Furthermore, the transient transformation of thalli with *Agrobacterium* carrying *intron-GUSPlus* revealed that Mp*npr-1^ge^* is more resistant to bacterial infection than Tak-1 (Figure 2F and 2G). These results indicate that MpNPR negatively regulates bacterial resistance in *M. polymorpha*, which implies that MpNPR and AtNPR1 play opposite roles in SA-mediated defense response.

### MpNPR is not a master regulator of SA-induced transcriptional reprogramming in M. polymorpha

In *A. thaliana*, the vast majority of SA-induced transcriptional reprogramming is governed by AtNPR1, AtNPR3, and AtNPR4 (Wang et al., 2006). Our results showing that there is potential for only a single SA receptor in *M. polymorpha*, MpNPR, would suggest that SA-induced transcriptional reprogramming will be lost in the Mp*npr* mutants. To investigate the SA-induced transcriptional response in *M. polymorpha* and the contributions of MpNPR to the response, 2-week-old Tak-1 and Mp*npr-1^ge^* plants were treated with 1 mM SA or Mock for 2, 6, and 12 hours and subjected to RNA-seq analysis (Figure S6A). The number of differentially expressed genes (DEGs, Mock vs SA, |Log_2_FC| ≥ 1, adjusted *p* < 0.05) in SA-treated Tak-1 peaked at 6 hours post-incubation (hpi) (Figure 3B), and thus we selected a 6 hpi dataset for further analyses. Strikingly, the overall transcriptional response pattern in Mp*npr-1^ge^* was found to be very similar to the pattern in Tak-1 (Figure 3A). Similar results were obtained when 7-day-old gemmae, which were grown in a liquid medium, were treated with SA and BTH (Figure S7). This indicates that the NPR protein is not a master regulator of SA-induced transcriptional reprogramming in *M. polymorpha*. Since the transcriptional response was rather pronounced in Mp*npr-1^ge^* compared to Tak-1 (Figure 3B), MpNPR may rather function as a repressor of SA-inducible transcription factors, which is in agreement with the SA hypersensitivity and the bacterial pathogen resistance phenotypes of the Mp*npr* mutants (Figure 2). Moreover, this implies that other, as yet unknown, molecules likely function as SA receptors in *M. polymorpha*.

**Figure 3.**
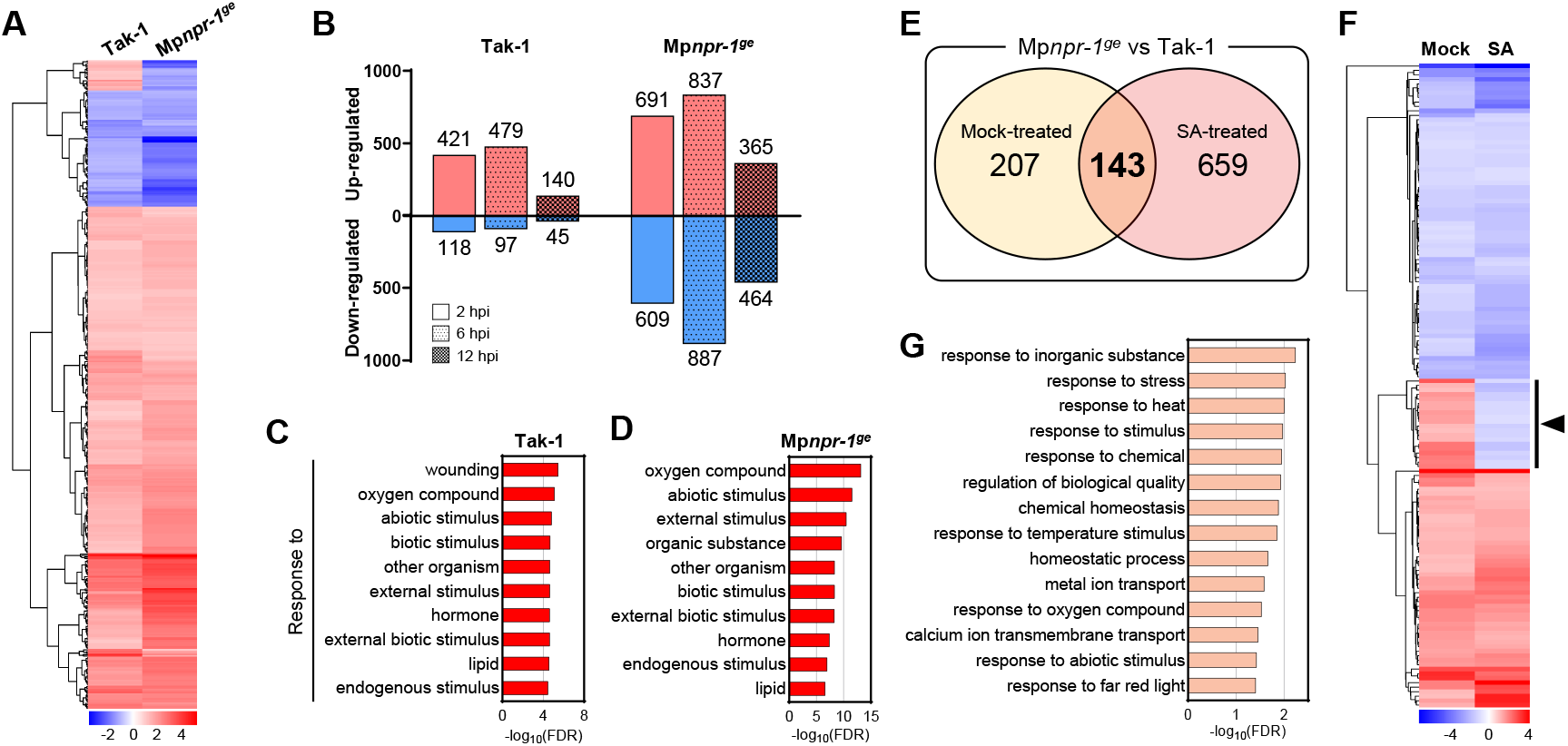
RNA-Seq analysis comparing SA-induced transcriptional reprogramming in Tak-1 and Mp*npr-1^ge^*. (A) Heatmap of 425 SA-induced DEGs (|log2FC| ≥ 1, adjusted *p* < 0.05, 6 hpi) identified in Tak-1 and correlated genes in Mp*npr-1^ge^*. (B) Number of SA-induced DEGs (|log2FC| ≥ 1, adjusted *p* < 0.05) in Tak-1 and Mp*npr-1^ge^* for 2, 6, and 12 hpi. (C, D) GO enrichment analyses using SA-induced DEGs in Tak-1 (C) and Mp*npr-1^ge^* (D). Top 10 GO terms beginning with ‘response to’ are shown (FDR < 0.001). (E) Venn diagram showing numbers of DEGs induced by Mp*npr-1^ge^* in each treatment (|log2FC| ≥ 1, adjusted *p* < 0.05, 6 hpi). (F) Heatmap of 143 DEGs identified in E, showing MpNPR-dependent and SA-irresponsive DEGs except for 20 genes (arrowhead). (G) GO enrichment analysis using 123 DEGs identified in F.

In Tak-1, 479 genes were up-regulated and 97 genes were down-regulated upon SA treatment (Figure 3B). Gene Ontology (GO) analysis showed significant enrichment of defense-related GO terms, including ‘*response to wounding*’ and ‘*response to other organism*’, and also GO terms related to responses to various stimuli in up- and down-regulated DEGs (Figure 3C). In Mp*npr-1^ge^*, 837 genes were up-regulated and 887 genes were down-regulated upon SA treatment (Figure 3B). Subsequent GO analysis revealed that GO terms ‘*response to wounding*’ is not enriched from the DEGs in Mp*npr-1^ge^* (Figure 3D). The expression of some Mp*PR* genes (Carella et al., 2019) was induced by SA treatment, and this up-regulation was slightly enhanced in Mp*npr-1^ge^* compared to Tak-1 (Figure S6C). These results imply that SA is involved in the defense response and that MpNPR contributes to the SA-dependent wounding-related response.

### MpNPR is involved in the heat and far-red light responses

To explore the biological processes that are mediated by NPR but are independent of the SA response in *M. polymorpha*, we compared the transcriptomes of Mp*npr-1^ge^* and Tak-1 in response to each treatment. Under Mock- and SA-treated conditions, 350 and 802 genes were identified as DEGs (Mp*npr-1^ge^* vs Tak-1, |Log_2_FC| ≥ 1, adjusted *p* < 0.05), respectively (Figure 3E). Among the DEGs, 143 genes were shared in both Mock- and SA-treated conditions (Figure 3E and 3F). In this subset, 20 genes showed significant changes in their expression patterns upon SA treatment (Figure 3F), and thus, we defined 123 genes as MpNPR-dependent and SA-nonresponsive genes. Subsequent GO analysis revealed that the biological processes ‘*response to heat*’, ‘*response to temperature stimulus*’, and ‘*response to far red light*’ are over-represented in these 123 genes (Figure 3G).

To examine whether MpNPR is involved in response to heat, we grew plants at high ambient temperatures, which was previously shown to induce folding and bending of *M. polymorpha* thalli as a thermomorphogenic response (Ludwig et al., 2021). As reported, enhanced three-dimensional (3D) growth of Tak-1 thalli was observed, resulting in reduced projected thallus area, when plants were grown at 30 °C (Figure 4A). The Mp*npr* mutants displayed enhanced folding and bending with smaller projected thallus area compared to Tak-1 (Figure 4A), indicating that MpNPR negatively regulates thermomorphogenesis in *M. polymorpha*. Then, we asked whether AtNPR1 also plays a role in thermomorphogenesis in *A. thaliana*. Strikingly, we found that the At*npr1-1* mutant and the At*npr1-1 npr3-2 npr4-2* triple mutant display enhanced hypocotyl elongation compared to a wild type Col-0 and the At*npr3-2 npr4-2* double mutant at 28 °C but not at 22 °C (Figure 4B). The *pif4-101* mutant was used as a control (Figure 4B). These results indicate that both MpNPR and AtNPR1 function as negative regulators of thermomorphogenesis.

**Figure 4.**
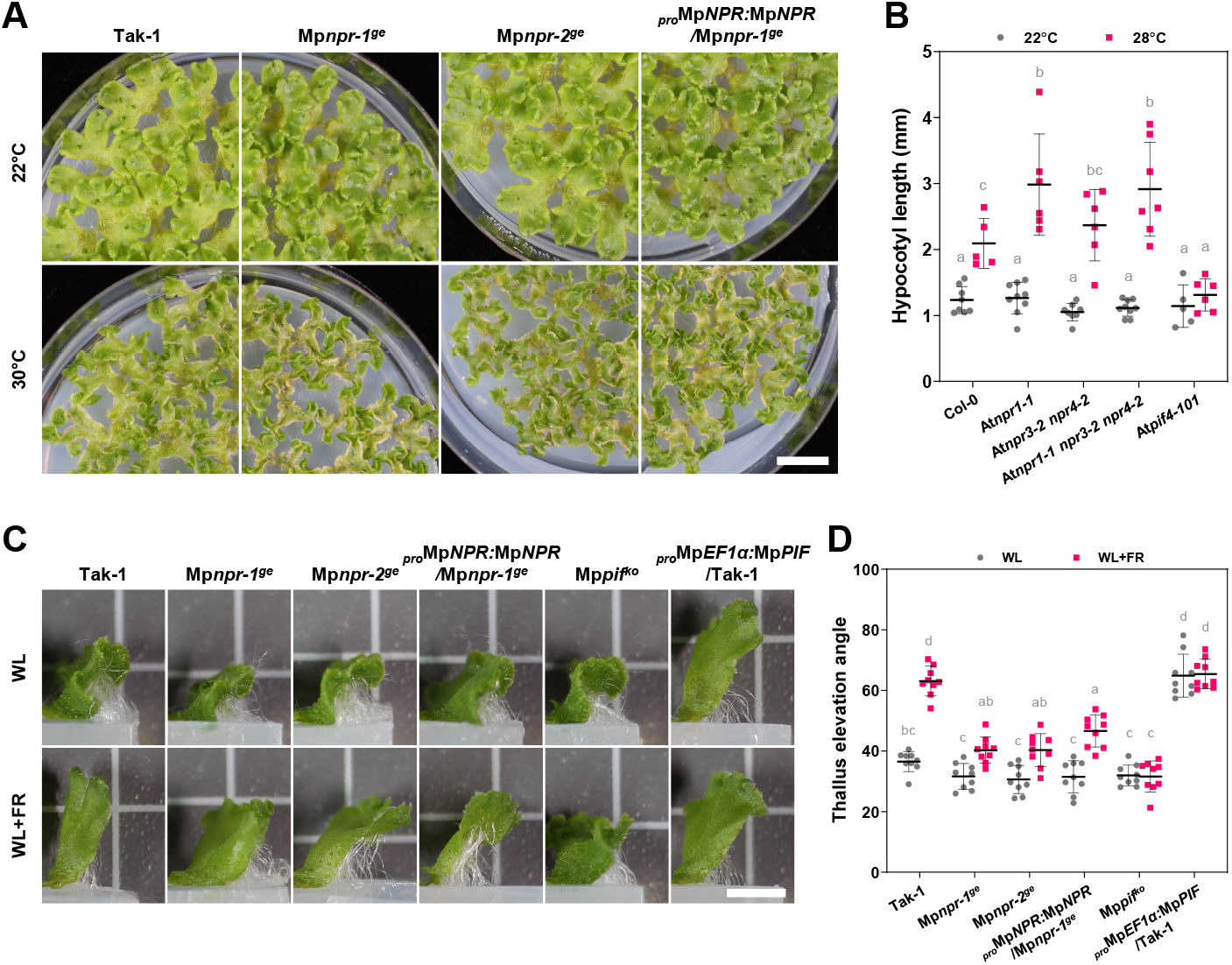
Putative roles of MpNPR in thermomorphogenesis and shade avoidance syndrome in *M. polymorpha*. (A) Enhanced thermomorphogenesis in Mp*npr* mutants compared to Tak-1 at 30 °C. Twenty-day-old thalli are shown. Scale bar, 1 cm. (B) Hypocotyl length of At*npr* mutants grown at 22 and 28 °C for 6 days (n ≥ 6). (C) Representative pictures of *M. polymorpha* thalli viewed from the side. Scale bar, 0.5 cm. (D) Thallus elevation angles quantified from 2-week-old *M. polymorpha* grown with or without far-red light (n = 9). Different letters represent statistically significant differences (Tukey’s test; *p* < 0.01). Error bars indicate SD.

We further investigated the possible contribution of MpNPR to far-red (FR) responses in *M. polymorpha*. When plants were grown in a condition supplemented with FR light, enhanced upward growth of thalli was observed, similar to shade avoidance responses in angiosperms (Figure 4C and 4D) (Casal et al., 1986; Franklin and Whitelam, 2005; Ballaré, 1999). The Mp*PIF* knockout and overexpressing plants were used as controls (Figure 4C and 4D). We found that Mp*npr* mutants display reduced upward growth compared to Tak-1 under the FR condition (Figure 4C and 4D). The complemented allele displayed a trend toward rescuing the phenotype but was not statistically significant (Figure 4D), which can be due to insufficient expression of MpNPR in this plant compared to Tak-1. This observation supports the hypothesis that MpNPR is involved in FR responses in *M. polymorpha*.

### Molecular properties are conserved between MpNPR and AtNPR1

Taking an inter-species complementation approach, we next determined whether the molecular properties of NPR proteins from *M. polymorpha* and *A. thaliana* are conserved. Expression of At*NPR1-mCitrine* under the Mp*EF1*α promoter in Mp*npr-1^ge^* rescued the SA-hypersensitivity phenotype (Figure 5A and 5B). Moreover, *_pro_*Mp*EF1*α:At*NPR1-mCitrine/*Mp*npr-1^ge^*plants displayed reduced SA sensitivity compared to Tak-1, which is probably due to overaccumulation of AtNPR1 (Figure 5A and 5B). Conversely, we expressed MpNPR under the cauliflower mosaic virus (CaMV) 35S promoter in the At*npr1-1* mutant. Expression of Mp*NPR* and At*NPR1* rescued the SA hypersensitive phenotype (Figure 5C). Line #4, which failed to express Mp*NPR*, was used as a control (Figure 5C). The At*npr1-1* mutant possesses the GUS reporter, *BGL2::GUS*, that is responsive to SA accumulation (Cao et al., 1994). In plants treated with SA, GUS-staining was observed only in the complementation lines (Figure 5D). Taken together, these results indicate that protein functions of MpNPR and AtNPR1 have been conserved to some degree during evolution. Furthermore, these results suggest that NPRs are associated with different components or pathways, which lead to distinct outputs, in different plant species.

## Discussion

Our phylogenetic analysis of 194 NPR proteins from 68 land plant species revealed that NPR proteins from angiosperms could be classified into two paralogous clades, NPR1/2 and NPR3/4, presumably diverged from an ancestral NPR (Figure 1). Two *NPR* genes were found in the genome of *Amborella trichopoda*, the single extant species of the sister lineage to all other angiosperms, and these genes, AmTr_v1.0_scaffold00036.47 and AmTr_v1.0_scaffold00032.72, were classified into the NPR1/2 and NPR3/4 clades, respectively (Dataset S1). Meanwhile, NPR proteins from gymnosperms formed a single cluster related to neither clade (Figure 1). This suggests that gene duplication of NPR led to the emergence of the two paralogous clades, seemingly associated with the epsilon whole-genome duplication (WGD) (Albert et al., 2013). Interestingly, the analysis showed that the AtNPR1/AtNPR2 and AtNPR3/AtNPR4 pairs most likely arose from a recent Brassicaceae-specific WGD event (Franzke et al., 2011; Ren et al., 2018), suggesting their functional redundancy in each clade. Indeed, AtNPR3 and AtNPR4 function redundantly, and AtNPR2, but not AtNPR3 and AtNPR4, can complement At*npr1-1* mutant phenotypes (Castelló et al., 2018; Zhang et al., 2006). Moreover, rice OsNPR1/NH1 and cacao TcNPR1, which belong to the NPR1/2 clade, could complement At*npr1* mutant phenotypes, and overexpression of OsNPR1/NH1 in rice resulted in enhanced resistance against the bacterial pathogen *Xanthomonas oryzae* pv. *oryzae* (Yuan et al., 2007). This suggests functional conservation among NPR proteins in the NPR1/2 clade. Similarly, cacao TcNPR3, which belongs to the NPR3/4 clade, could complement At*npr3-3* mutant phenotypes, and CRISPR/Cas9-based mutagenesis of *TcNPR3* in cacao resulted in enhanced resistance against the cacao pathogen *Phytophthora tropicalis*, suggesting functionally conserved roles for NPR proteins in the NPR3/4 clade as negative regulators of defense responses (Fister et al., 2018; Shi et al., 2013). However, there is a contradictory report which showed that overexpression of *Nicotiana glutinosa* NgNPR3, an AtNPR3 homolog, in *Nicotiana tabacum* cv. Samsun enhanced resistance to *Alternaria alternata*, *Pseudomonas solanacearum*, and potato virus Y (Zhang et al., 2010b). Nevertheless, assuming that NPR proteins from the paralogous clades NPR1/2 and NPR3/4 have acquired opposed or different functions, an obvious question arises of whether and how NPR proteins in the non-seed plants clade, which can be remnants of NPR in the common ancestor of land plants, play roles in SA signaling and defense responses. A single study reported that *P. patens* PpNPR1 partially complements phenotypes of At*npr1-1* but not of At*npr3 npr4* (Peng et al., 2017). Together with our finding that AtNPR1 and MpNPR are interchangeable in the range of evaluated phonotypes, it is likely that the molecular properties of non-seed plant NPR proteins and NPR proteins in NPR1/2 clade have been relatively well-conserved during evolution. If so, a remaining question is whether the molecular properties of NPR proteins in the NPR3/4 clade are distinct from non-seed plant NPR proteins. Further genetic studies in many other plant species and inter-species complementation assays will enhance our understanding of the functional conservation and diversification of NPR proteins.

There is compelling evidence that AtNPR1 is the master regulator of SA- or BTH-induced transcriptional reprogramming in *A. thaliana*. However, there is as yet no hint that other Brassicaceae NPR1-like proteins or NPR proteins in the NPR1/2 clade also function as master regulators in their respective species. In order to genetically engineer *NPR* genes to manipulate disease resistance, it is crucial to understand the molecular functions of NPR proteins in other plant species. Further, it remains unclear how AtNPR3 and AtNPR4 have evolved to become transcriptional co-repressors with opposite roles to AtNPR1. One model that has been proposed is that the EAR motif located in the NPR1-like-C domain of AtNPR3 and AtNPR4 differentiates them from AtNPR1, which lacks the EAR motif (Ding et al., 2018). We found that the EAR motif found in AtNPR3 and AtNPR4 is perfectly conserved in all Brassicaceae NPR2-like proteins (Figure S1), which suggests that Brassicaceae NPR2-like proteins can rather function as transcriptional co-repressors instead of co- activators like AtNPR1. However, as mentioned above, AtNPR2 was shown to complement At*npr1-1* mutant phenotypes. Conservation of the EAR motif in Brassicaceae NPR2-like proteins suggested to us that AtNPR1 or Brassicaceae NPR1-like proteins have presumably lost the EAR motif during evolution. To evaluate this hypothesis, we expanded our analysis of functional protein sequences to all 194 NPR proteins. As expected, we found that the EAR motif is widely conserved in NPR proteins from gymnosperms and all three major clades except for Brassicaceae NPR1-like subclade (Figure S1). It is formally possible that other NPR proteins have acquired the EAR motif independently during evolution, but we suggest that the more reasonable assumption is that Brassicaceae NPR1-like proteins have lost the EAR motif after a Brassicaceae-specific WGD event. As mentioned above, rice OsNPR1/NH1, cacao TcNPR1, and *P. patens* PpNPR1 could complement phenotypes in At*npr1* mutants (Peng et al., 2017; Shi et al., 2010; Yuan et al., 2007), and these three NPR proteins carry the EAR motif. These observations indicate that the EAR motif is not by itself sufficient for the co-repressor or co-activator function of NPR proteins. One plausible explanation could be that the EAR motif is controlled by phosphorylation. We found that SP or TP sites in most NPR proteins of dicots in the NPR1/2 clade were substituted with non-phosphorylable hydrophobic residues (A/V/L/I/M) (Figure S1). Although we have no direct evidence that these sites are targeted by phosphorylation, these clade-specific mutations may further explain how the EAR motif determines protein function.

The remarkable conservation of the Arg residue among NPR proteins, which is required for SA binding in AtNPR1, AtNPR3, and AtNPR4 (Ding et al., 2018; Fu et al., 2012; Wu et al., 2012), seemingly suggests that NPR proteins generally function as SA receptors in plants. However, unexpected conservation of the EAR motif made it difficult for us to predict how NPR proteins coordinate the SA response in non-Brassicaceae plant species. Rice, *O. sativa*, has three NPR proteins, OsNPR1/OsNH1, OsNPR2/OsNH2, and OsNPR3 (Yuan et al., 2007), of which OsNPR1/OsNH1 belongs to the NPR1/2 clade and OsNPR2/OsNH2 and OsNPR3 belong to the NPR3/4 clade (Figure S1). Overexpression of these three genes in rice suggested that only OsNPR1/OsNH1 functions as a positive regulator of defense responses (Yuan et al., 2007), that is, OsNPR1/OsNH1 but not OsNPR2/OsNH2 or OsNPR3, is a transcriptional co-activator of SA signaling. Accordingly, RNAi-mediated knockdown of Os*NPR1* in rice resulted in impairment of BTH-induced transcriptional reprogramming, but the observed impairment was not as striking as in At*npr1* mutants (Sugano et al., 2010). It is important to generate and analyze Os*npr1* knockout plants, but there is a possibility that release of transcriptional repression by OsNPR2/OsNH2 and OsNPR3 makes a large contribution to the SA response. Analysis of a Os*npr1 npr2 npr3* triple mutant would clarify the contribution of NPR to SA signaling in rice. We found that two NPR proteins in *P. patens* carry the EAR motif but that the single NPR in *M. polymorpha* lacks this motif (Figure S1). The observation that the expression of MpNPR or PpNPR1 in At*npr1-1* can restore the SA response suggests that bryophyte NPR proteins can function as SA receptors and transcriptional co-activators in *A. thaliana*, irrespective of the conservation of EAR motif.

It is striking that mutation of the single *NPR* gene in *M. polymorpha*, Mp*NPR*, did not result in diminished SA-induced transcriptional responses, although MpNPR can function as a transcriptional co-activator in *A. thaliana*. Our transcriptomic analysis suggests that MpNPR may function as a transcriptional co-repressor in *M. polymorpha* (Figure 3 and S6C). In a similar fashion, the Mp*npr* mutants displayed enhanced resistance against bacterial pathogens (Figure 2E to 2G). This is an interesting observation because it implies that NPR is not important for *M. polymorpha* resistance against biotrophic or hemi-biotrophic pathogens. Together with the evidence that hornworts have lost the *NPR* gene (Li et al., 2020), NPR may play a less important role in disease resistance in bryophytes. It would be instructive to investigate the functions of NPR in mosses in the future, but we speculate that plants have acquired the *NPR* gene during terrestrialization initially for purposes other than SA-mediated immunity. Besides, it is important to understand why loss of MpNPR or AtNPR1 leads to distinct phenotypes in each species, even though MpNPR and AtNPR1 are exchangeable at some level. Further identification of components that interact with NPRs would help to understand molecular mechanisms behind these intriguing differences.

Our analyses highlighted that Brassicaceae species have evolved a unique NPR repertoire. Marked changes to amino acid sequence were observed to have accumulated in Brassicaceae NPR1-like proteins (Figure 1B and S1). Although we still do not know how these mutations contribute to the molecular functions and physiological roles of Brassicaceae NPR1-like proteins, this finding is rather surprising considering the importance of AtNPR1 in SA signaling and the immune system in *A. thaliana*. Among four Brassicaceae-specific subclades, further lineage-specific expansion of NPR1 was observed (Figure S3A). This may suggest that Brassicaceae *NPR1-like* genes are experiencing selective pressures. In this respect, AtNPR1 was shown to be targeted by the *P. syringae* type III effector AvrPtoB and *Phytophthora capsici* effector RxLR48 (Chen et al., 2017; Li et al., 2019), further confirming the significance of AtNPR1 in immunity in *A. thaliana*. Effector targets are occasionally integrated into NLR proteins as decoys to counteract pathogen infection (Sarris et al., 2016). Strikingly, we found that one of the *Boechera stricta* NPR1 homologs, Bostr.29223s0069.1, is integrated into the C-terminus of TIR-NLR (Figure S3C). Interestingly, BLASTP comparison of this TIR-NLR against the *A. thaliana* database returned SNC1 (suppressor of *npr1-1*, constitutive 1) as a best hit (Zhang et al., 2003). The expression of the TIR-NLR-NPR as a full-length protein needs to be confirmed, but this information seemingly suggests that Brassicaceae NPR1-like proteins generally play a key role in SA-mediated immunity in Brassicaceae species.

What then might have been the ancestral functions of NPR or NPR-associated pathways in the common ancestor of land plants? Our transcriptomic analysis suggested potential roles of MpNPR in heat and FR signaling in *M. polymorpha*. We indeed found that the Mp*npr* mutants display enhanced thermomorphogenesis and reduced FR-induced shade avoidance (Figure 4A, 4C, and 4D). Importantly, we revealed that At*npr1-1* also displays enhanced thermomorphogenesis (Figure 4B), implying that a function of NPR in thermomorphogenesis is conserved in the only two sister lineages of land plants, bryophytes and tracheophytes, and is thus ancient. At*npr1-1* was also reported to display reduced shade avoidance (Nozue et al., 2018). Considering that adaptation to fluctuating temperatures was crucial for plant terrestrialization, it is attractive to hypothesize that plants initially acquired *NPR* genes to deal with varying abiotic components, including temperature and light quality, rather than potentially detrimental microbes. In comparison with the NPR family, the BOP family seems to have undergone less expansion during land plant evolution. The genome of *A. trichopoda* encodes a single *BOP* gene (Albert et al., 2013), suggesting that *BOP* was not duplicated at the epsilon WGD event. In this regard, the two BOP proteins encoded in the *A. thaliana* genome function redundantly (Hepworth et al., 2005; Norberg et al., 2005; Xu et al., 2010). Nevertheless, analyzed plant species encode between zero and five copies of BOP, suggesting lineage-specific expansion and loss of BOP. The function of *BOP* genes in bryophytes is not yet known, and disruption of all three *BOP* genes in the moss *P. patens* did not cause any observable developmental phenotypes (Hata et al., 2019). It is intriguing that *Marchantia* liverworts have kept the *NPR* gene but lost the *BOP* gene, while *Anthoceros* hornworts have lost the *NPR* gene but kept the *BOP* gene (Bowman et al., 2017; Li et al., 2020). This may imply that *BOP* and *NPR* genes have shared functions that were required for the survival of liverwort and hornwort lineages during evolution. In *A. thaliana*, two BOPs have been shown to play a role in thermomorphogenesis (Zhang et al., 2017), suggesting that AtNPR1 and AtBOP1/2 have shared functions in thermomorphogenesis. Functional analysis of *BOP* genes in hornworts and sophisticated inter-species and intra-species complementation analysis would shed light on the ancestral shared functions of NPR and BOP proteins or of a hypothetical ancestral protein that diverged into NPR and BOP.

## Methods

### Key resources table

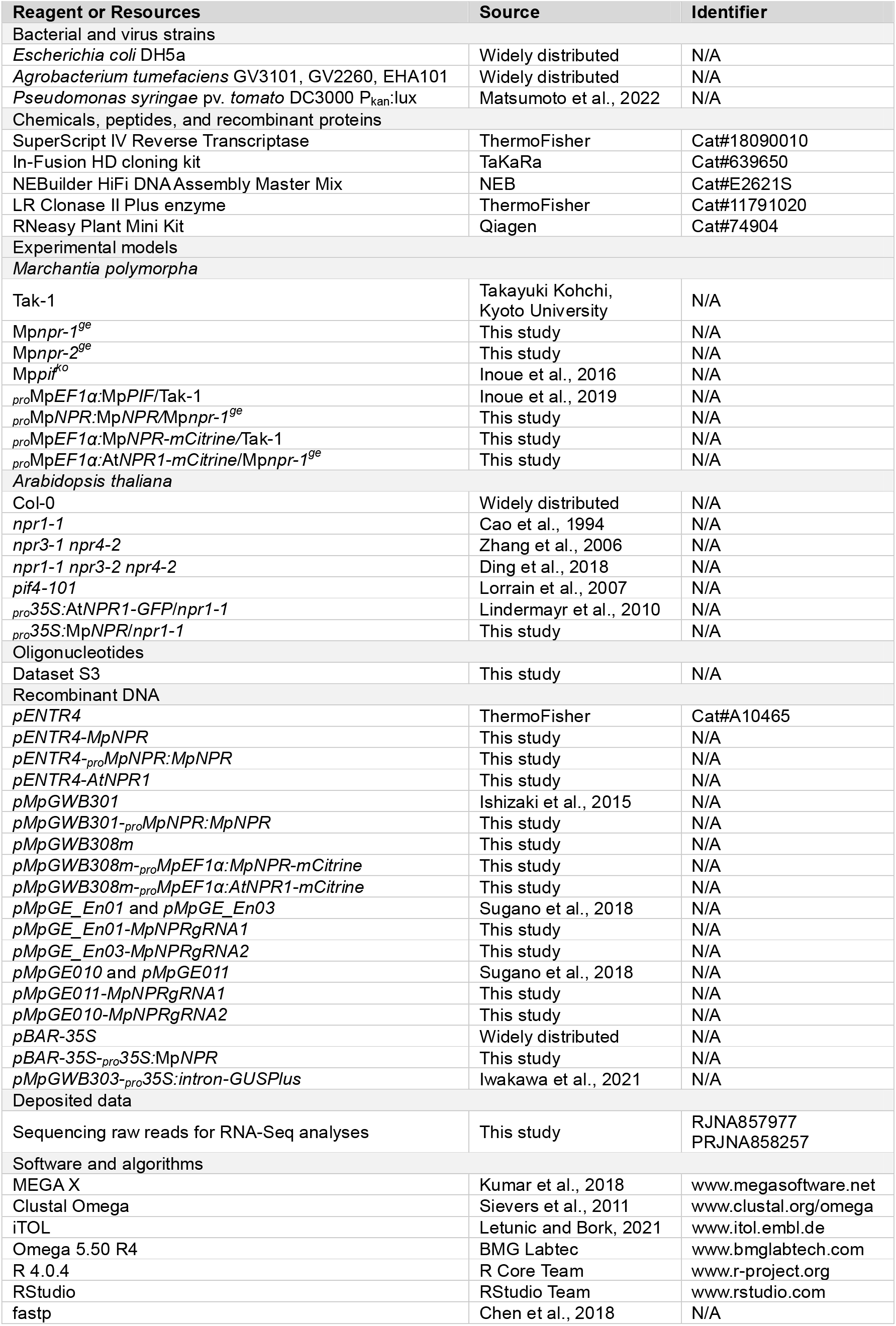
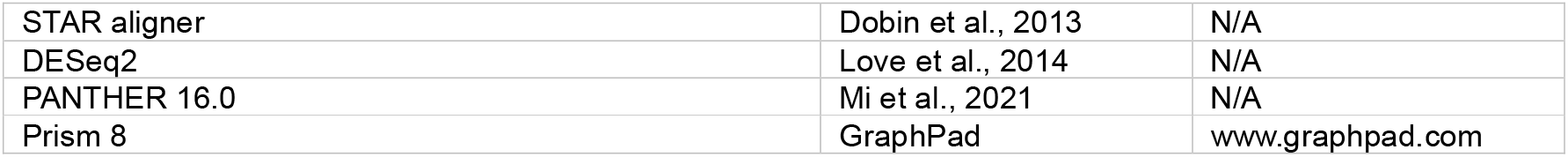

### Experimental model and subject details

*Marchantia polymorpha* accession Tak-1 (Ishizaki et al., 2008) was used as a wild type throughout this study. For cultivation, gemmae were grown on half-strength Gamborg’s B5 (GB5) basal media containing 1% agar at 22°C under continuous white light (60 to 70 μmol m^-2^ s^-1^). Sodium salicylate (Sigma-Aldrich) was added to the media for SA-supplemented conditions to achieve desired concentrations. For far-red (FR) light irradiances, gemmae were cultivated in the growth chamber (Percival Scientific), maintaining continuous white light (400 to 700 nm, 65 μmol m^-2^ s^-1^) equipped with FR LEDs (700 to 780 nm, 65 μmol m^-2^ s^-1^) at 22°C. Light properties were measured by LI-250A light meter (LI-COR) equipped with LI-190R quantum sensor (LI-COR) or SKR110 R/FR sensor (Skye instruments).

*Arabidopsis thaliana* ecotype Col-0 was used as a wild type in this study. For cultivation of *A. thaliana*, seeds were surface-sterilized by vapor gas (Sodium hypochlorite containing 3% [v/v] HCl) for 4 hours and placed on half-strength Murashige and Skoog (MS) media containing 1% agar. Seeds were then incubated at 4°C in the dark for 2 days before germination. Germinated seedlings were cultivated at 22°C under continuous white light (60 to 70 μmol m^-2^ s^-1^). For the SA sensitivity test, sodium salicylate (Sigma-Aldrich) was added to the media to achieve desired concentrations.

### Cloning and Plasmid Construction

To generate Mp*npr-1^ge^* and *Mpnpr-2^ge^*using CRISPR/Cas9 system, the first exon of Mp*NPR* was targeted (Figure S4A). The double-strand oligonucleotides containing the target sequences were synthesized and cloned into *pMpGE_En01* or *pMpGE_En03* using In-Fusion HD cloning kit (TaKaRa) and BsaI restriction sites in *pMpGE_En03*, respectively. Generated entry clones were subsequently used for the recombination into destination vectors *pMpGE011* and *pMpGE010* by Gateway LR clonase (ThermoFisher) as manufacturer’s instruction.

For complementation of *M. polymorpha* Mp*npr* mutants, the full-length CDS of Mp*NPR* (Mp1g02380) and At*NPR1* (At1g64280) were first amplified from cDNA of *M. polymorpha* Tak-1 and *A. thaliana* Col-0, respectively. For the MpNPR promoter, 5.6 kb upstream of ATG was amplified from Tak-1 genomic DNA. The amplified fragments were assembled in *pENTR4* (ThermoFisher) using HiFi assembly master mix (NEB), followed by recombination into *pMpGWB301* or *pMpGWB308m* (modified version of *pMpGWB308* (Ishizaki et al., 2015), replacing Citrine to mCitrine) using Gateway LR clonase (ThermoFisher). For complementation of *A. thaliana* At*npr1-1* mutant, the full-length CDS of Mp*NPR* was amplified from cDNA of *M. polymorpha* ecotype BoGa and cloned into *pBAR-35S* using XmaI and XbaI restriction sites.

### *M. polymorpha* transformation

The resulting constructs were transformed into *Agrobacterium tumefaciens* GV3101, GV2260 or EHA101 by electroporation. Subsequent transformations were carried out as described before (Kubota et al., 2013). Obtained transformants were grown on half-strength GB5 basal media containing 1% agar supplemented with 100 μg/mL cefotaxime and 10 μg/mL hygromycin B or 0.5 μM chlorsulfuron to screen non-chimeric transgenic gemmae.

### GUS histochemical assay

10-day-old Mock- and 0.1 mM SA-grown *A. thaliana* seedlings were submerged in GUS staining solution consisting of 0.5 mg/mL X-Gluc (5-bromo-4- chloro-3-indolyl-beta-D-glucuronic acid), 0.1% Triton X-100, 10 mM EDTA, 0.5 mM potassium ferricyanide, 0.5 mM potassium ferrocyanide in 100 mM sodium phosphate buffer (pH 7.0), followed by vacuum infiltration for 20 min. After incubation in the dark at 37°C for overnight, leaves were destained by serial incubation in 30 %, 50 %, 70 % ethanol for a minimum of 1 hour each before observation.

### Phylogenetic analyses

Amino acid sequences of 317 homologs of MpNPR and AtBOP1 (Dataset S1) were aligned using Clustal Omega with default parameters. To assess the reliability of a phylogenetic tree, the alignment was further calculated using the Maximum Likelihood method with a bootstrap value of 1000 in MEGA software (Kumar et al., 2018).

### Phytohormones measurements

*M. polymorpha* thalli were grown on Mock and 0.1 mM SA for 14 days before homogenization. Phytohormones were extracted with methyl-tert-butyl ether (MTBE), reversed phase-separated using an ACQUITY UPLC system (Waters Corp.) and analysed by nanoelectrospray ionization (nanoESI) (TriVersa Nanomate, Advion BioSciences) coupled with an AB Sciex 4000 QTRAP tandem mass spectrometer (AB Sciex) employed in scheduled multiple reaction monitoring modes (Herrfurth and Feussner, 2020). For SA measurement, the following mass transition was included 137/93 [declustering potential (DP) −25 V, entrance potential (EP) −6 V, collision energy (CE) −20 V].

### Bioluminescence-based bacteria quantification

Bacterial quantification in infected thallus was carried out as described before (Matsumoto et al., 2022). Briefly, *M. polymorpha* were grown on autoclaved cellophane disc on half-strength GB5 media for 2 weeks. In the meantime, *Pto*-lux was cultivated in King’s B medium containing 30 μg/mL rifampicin to achieve 1.0 of OD_600_. The saturated bacteria culture was subsequently washed and resuspended in Milli-Q water to prepare bacteria suspension with 0.01 of OD_600_. Next, 2-week-old thalli were submerged in the bacteria suspension followed by vacuum for 5 min and incubated for 0 to 3 days on humid filter papers. After incubation, thallus discs (5 mm diameter) were punched from the basal and apical region using a sterile biopsy punch (pfm medical) and transferred to a 96-well plate (VWR). The bioluminescence was measured in the FLUOstar Omega plate reader (BMG Labtech).

### *Agrobacterium*-mediated transient transformation

Transient transformation was carried out as described before (Iwakawa et al., 2021). A single colony of *Agrobacterium tumefaciens* strain GV3101::pMP90 harboring *_pro_35S:intron-GUSPlus* was inoculated into Luria-Bertani medium containing spectinomycin antibiotics and cultured for 2 days at 28°C. Bacterial cells were collected from 1 ml culture by centrifugation, suspended in 5 ml of 0M51C medium (Ono et al., 1979; Takenaka et al., 2000) containing 2% sucrose and 100LµM 3,5-dimethoxy-4-hydroxyacetophenone (acetosyringone) and cultured for 5 hours at 28 °C. In parallel, Tak-1 and Mp*npr-1^ge^* gemmae were cultured on the agar plates for 14 days at 22 °C under continuous light. The bacterial culture was diluted with 0M51C medium containing 2 % sucrose and 100 µM acetosyringone to be at 0.02 of OD600, and then thalli were transferred into the 50 ml bacterial suspension. The thalli and *Agrobacterium* were co-cultured at 22 °C under continuous light for 3 days with shaking (120 rpm).

### RNA-Seq and data analysis

Total RNA was isolated from 2-week-old *M. polymorpha* Tak-1 and Mp*npr-1^ge^* treated with Mock or 1 mM SA for 2, 6, and 12 hours. For RNA-Seq analysis of 7- day-old gemmae grown in liquid half-strength GB5 medium with 0.1% (w/v) sucrose at 22 °C under continuous light with shaking (120 rpm), total RNA was isolated from *M. polymorpha* Tak-1 and Mp*npr-1^ge^*gemmae treated with Mock (water), 1 mM SA, Mock (DMSO), and 0.5 mM BTH for 24 hours. Library preparation and sequencing were performed by the Max Planck-Genome-center, Cologne (https://mpgc.mpipz.mpg.de), on the Illumina HiSeq 3000 platform. Raw reads were processed by fastp (Chen et al., 2018) for quality control and trimming. The *M. polymorpha* genomes and gene annotations (MpTak_v6.1) (Montgomery et al., 2020) were used for mapping reads and counting transcripts per gene in STAR aligner (Dobin et al., 2013). The genes with less than the average of 10 read counts were excluded, and the log2 fold difference of the gene expression between conditions was calculated in R package DESeq2 (Love et al., 2014). Genes with statistical significance (FDR adjusted p-value < 0.05) were selected for further analyses.

### Quantification and statistical analysis

Statistical details, including sample numbers, error bars, and p-value cut-offs, were shown in corresponding figure legends. Unless otherwise specified, all statistical significances were calculated using the R function pairwise.t.test with pooled SD. The Benjamini–Hochberg method was then used for correcting the multiple hypothesis testing. Linear regression and calculation of correlation coefficients were performed using the R function lm and cor, respectively.

### Data availability

Sequencing raw reads used in transcriptomic analyses in this study have been deposited under the accession BioProject PRJNA857977 and PRJNA858257.

## Acknowledgements

The authors are grateful to Takayuki Kohchi (Kyoto University, Japan) for providing pMpGWB and pMpGE vectors and Mp*pif^ko^*and *_pro_*Mp*EF1*α:Mp*PIF*/Tak-1 plants, Yuelin Zhang (The University of British Columbia, Canada) for providing At*npr1-1*, At*npr3-2 npr4-2*, and At*npr1-1 npr3-2 npr4-2* seeds, George Coupland (MPIPZ, Germany) for providing At*pif4* seeds, and Sabine Freitag (University of Göttingen, Germany) for expert technical assistance. We thank Neysan Donnelly (MPIPZ, Germany) for editing the manuscript.

## Author contributions

H.-W.J., H.I., and H.N. designed the research. H.I. performed phylogenetic and alignment analysis. H.I., S.N., and J.K. generated Mp*npr* mutants. H.-W.J. and H.I. generated Mp*npr-1^ge^* complementation and MpNPR overexpression lines. N.G. and S.Z. generated At*npr1-1* complementation lines. H.-W.J. and H.I. performed SA and BTH sensitivity assays. C.H. and I.F. quantified phytohormones. H.-W.J. and T.S. performed bioluminescence-based bacteria quantification. H.I. performed transient transformation assay. H.-W.J., H.I., and S.M. performed transcriptomic analyses. H.-W.J. performed all other experiments. H.-W.J., H.I., and H.N. wrote the manuscript. All authors corrected the manuscript.

## Supplemental information

**Dataset S1** NPR and BOP homologs used for phylogenetic analysis

**Dataset S2** Reference genomes of species used in BLASTP

**Dataset S3** Oligonucleotides used in this study

---

**Supplementary Figure S1.**
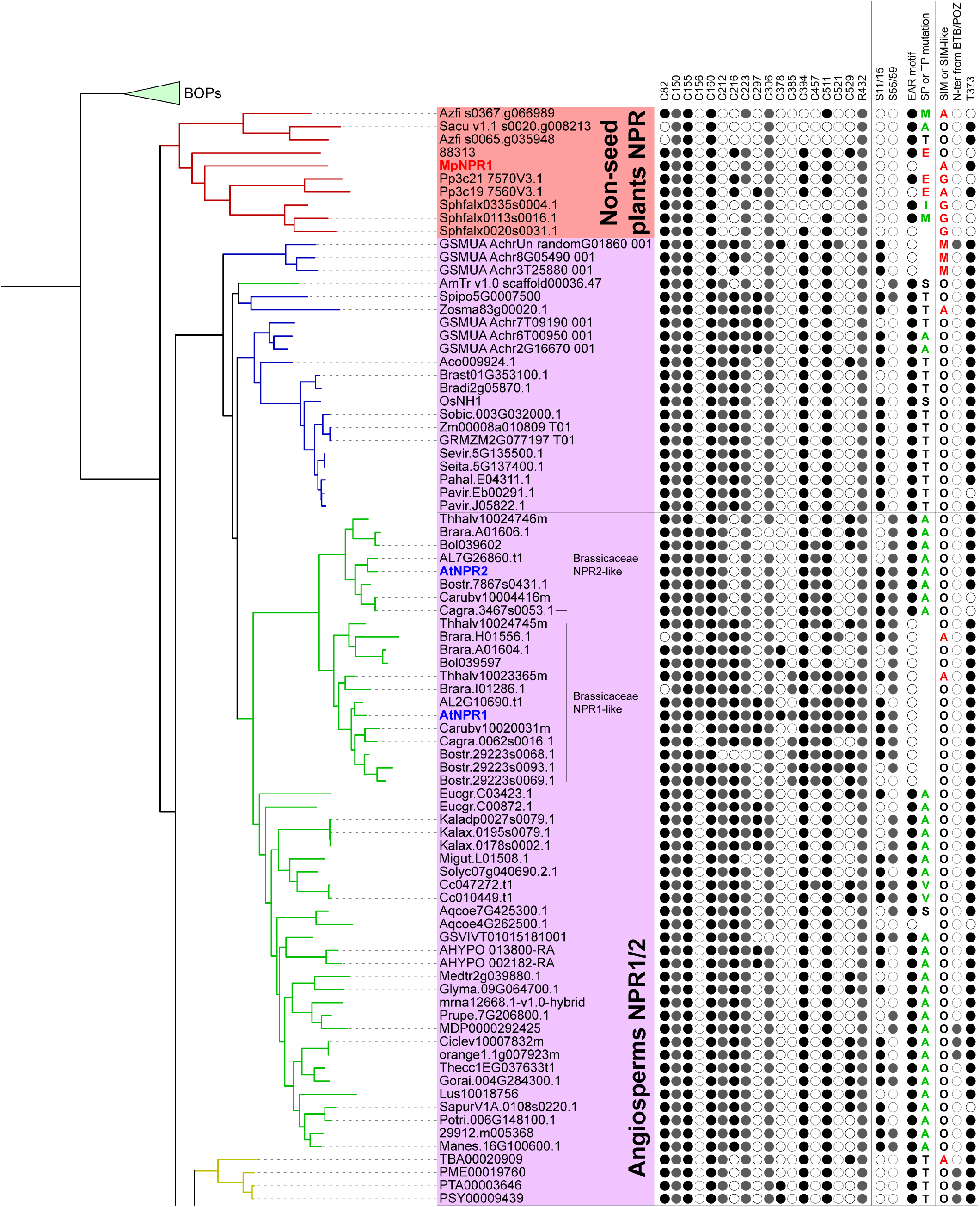
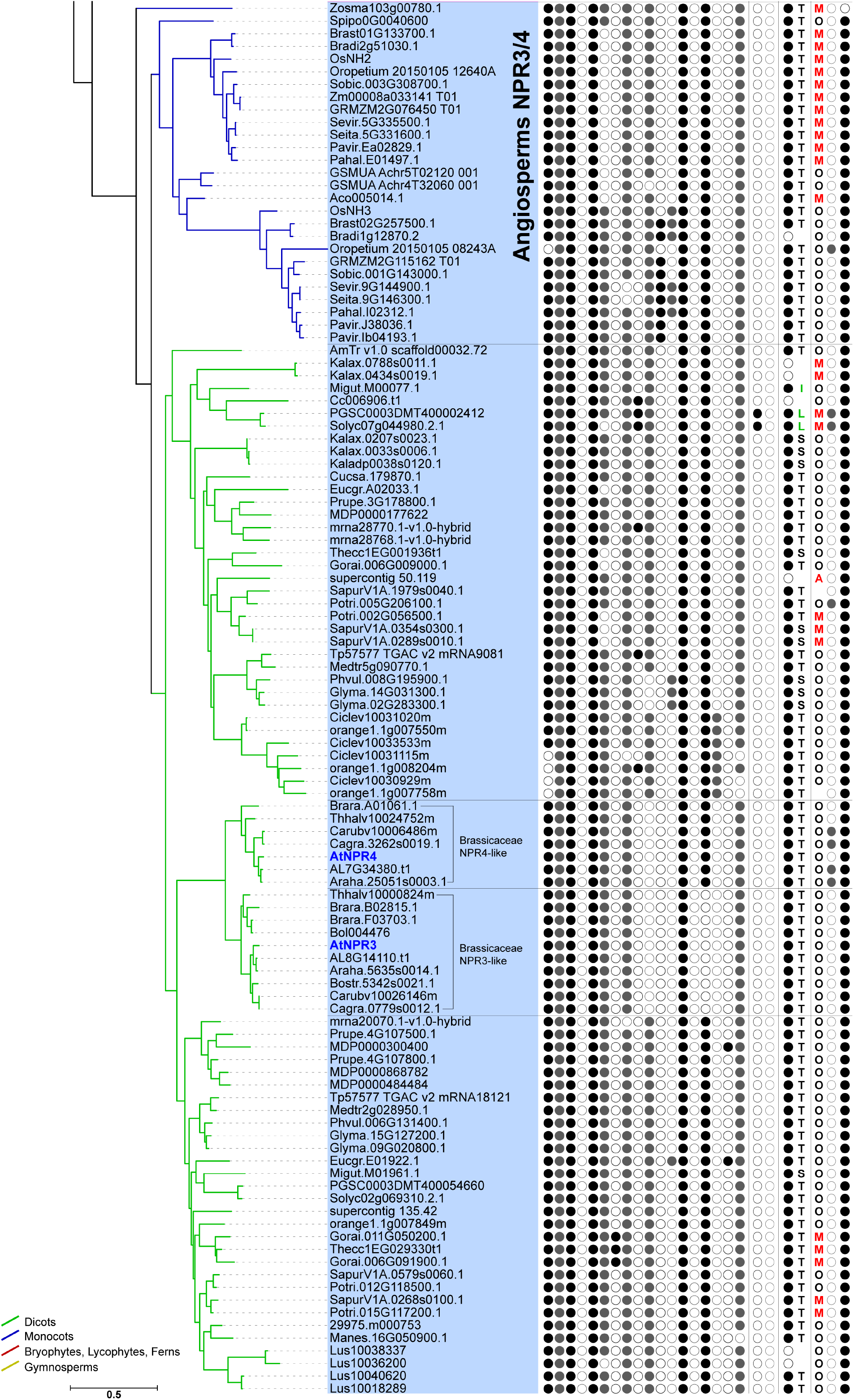
Phylogenetic tree showing the conservations of functional residues and motifs of 194 NPR proteins, related to Figure 1. Presence and absence of functional residues and motifs are shown by closed and open circle, respectively. In SP/TP mutations, substitutions with non-phosphorylatable hydrophobic residues (A/V/L/I/M) are colored in green.

**Supplementary Figure S2.**
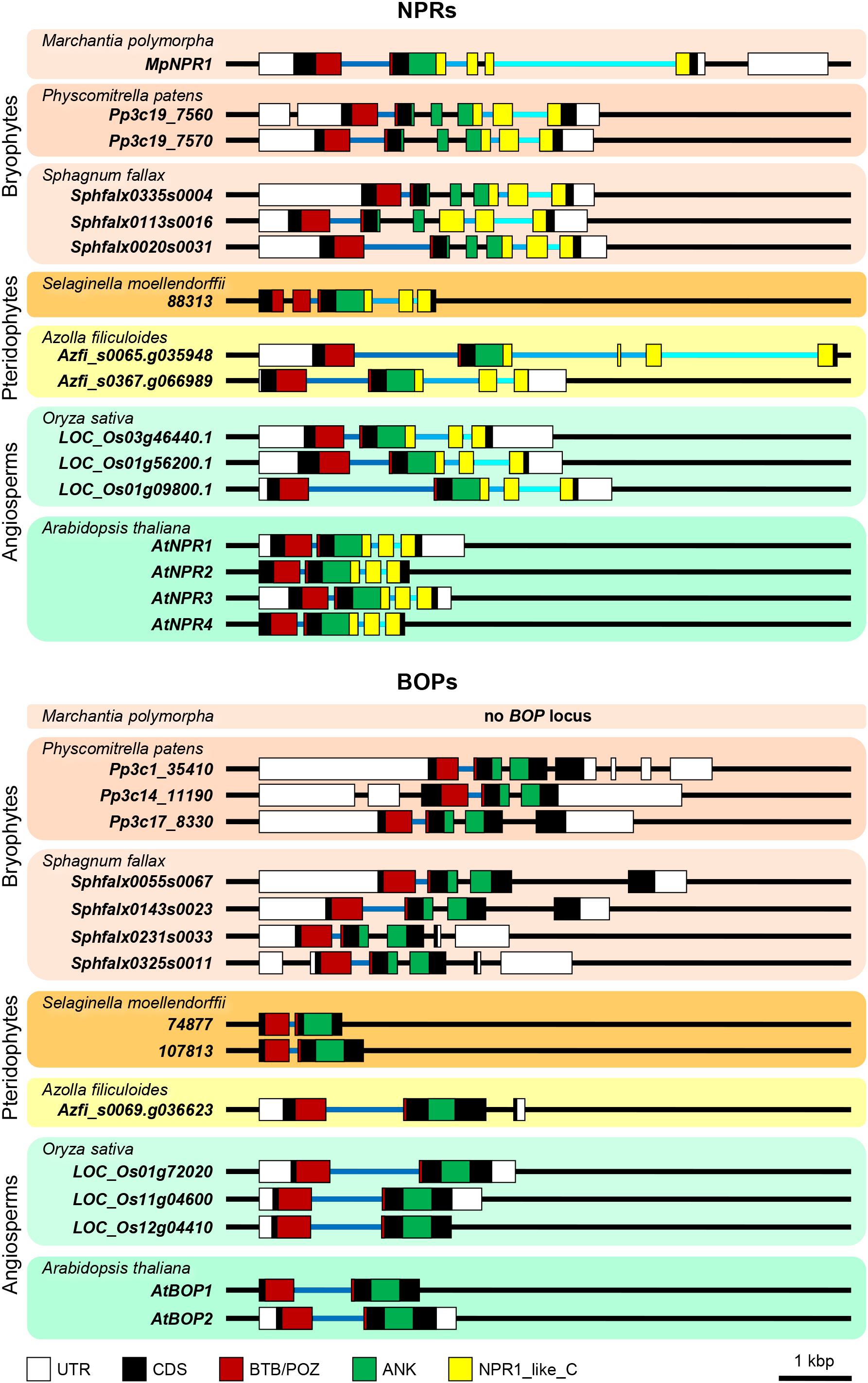
The exon-intron organization in the BTB/POZ-encoding regions are highly conserved between NPR and BOP families.

**Supplementary Figure S3.**
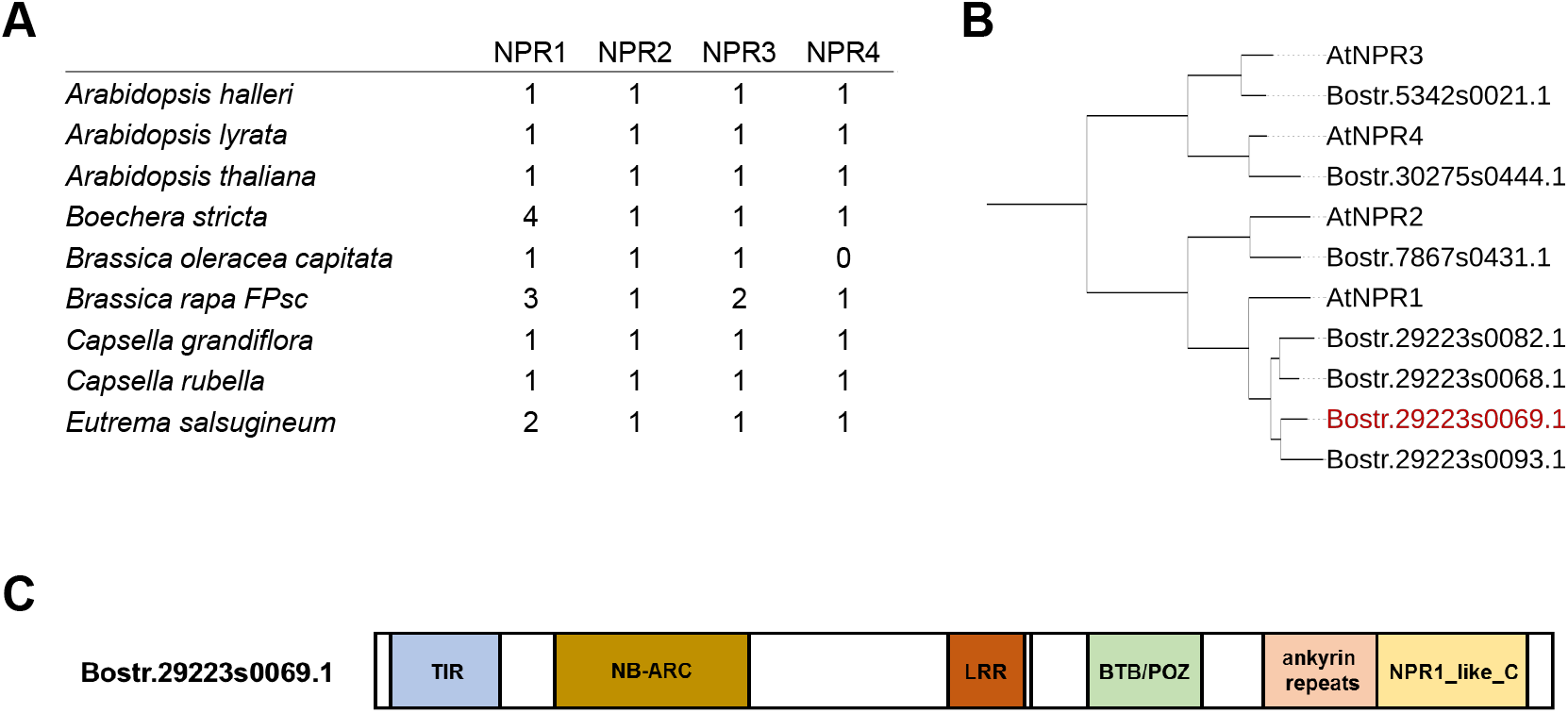
Gene duplications in Brassicaceae-specific NPR subclades and integration of *Boechera stricta* NPR1 homolog into TIR-NLR protein, related to Figure 1A. (A) Number of NPRs in Brassicaceae species suggesting the selective pressures or expansions of Brassicaceae NPR-like genes. (B) Phylogenetic analysis of *Boechera stricta* NPR homologs. (C) Schematic of domains showing that Bostr.29223s0069.1 is integrated into the C-terminus of TIR-NLR proteins in *Boechera stricta*.

**Supplementary Figure S4.**
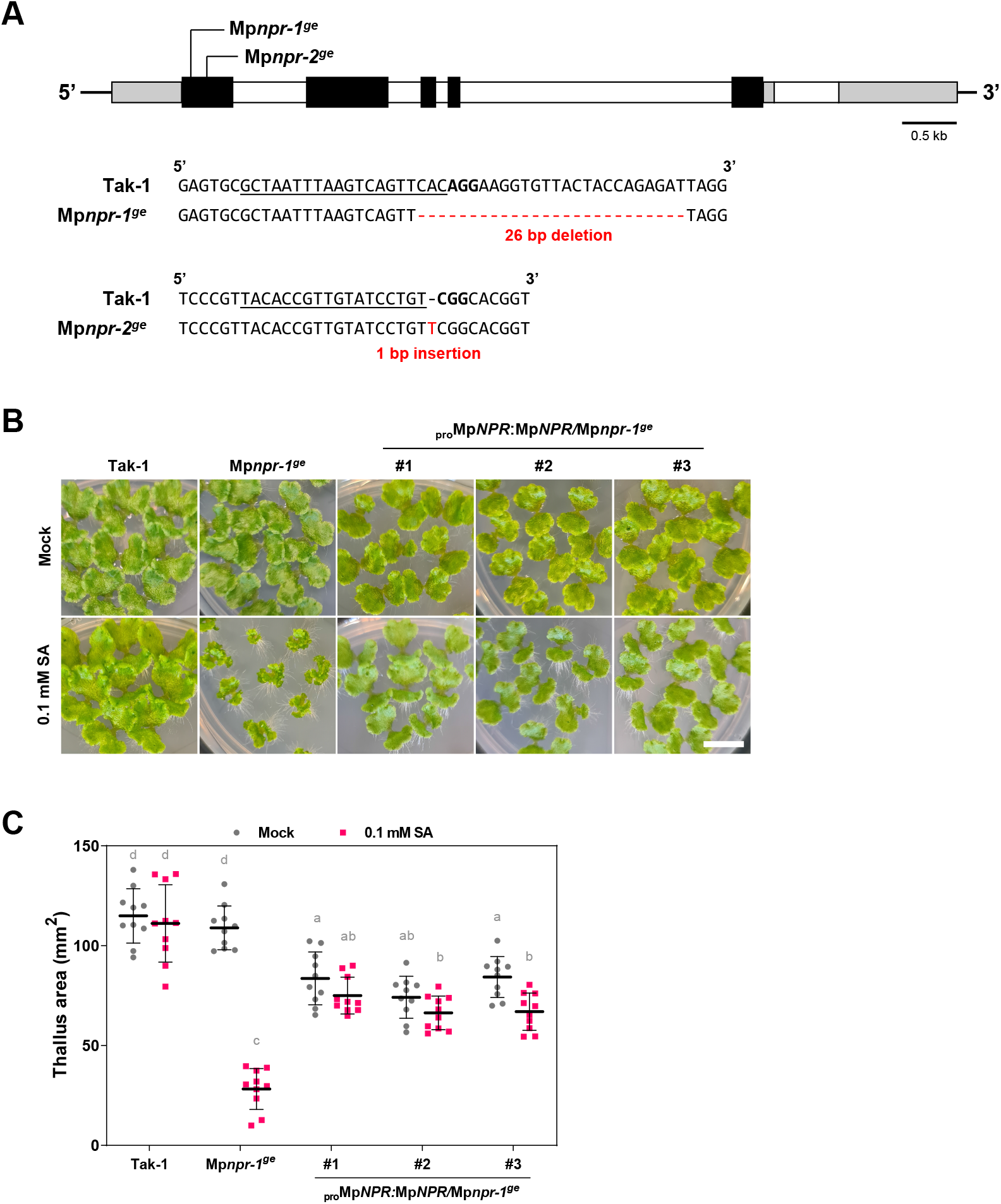
Generation of Mp*npr* mutants using CRISPR/Cas9 system and complementation of the SA-hypersensitivity phenotype in the mutants. (A) Schematic of sgRNA target sites in the first exon of Mp*NPR*. Black, white and gray boxes indicate exons, introns, and UTRs, respectively. Mp*npr-1^ge^* and Mp*npr-2^ge^* resulted in 26 bp deletion and 1 bp insertion, respectively. (B) Complementation of SA-hypersensitivity in individual transgenic lines expressing _pro_Mp*NPR:*Mp*NPR*. Two-week-old *M. polymorpha* grown on mock and 0.1 mM SA conditions are shown. Scale bar, 1 cm. (C) Quantification of thallus projected area shown in B. Different letters represent statistical significances (Tukey’s test; *p* < 0.01). Error bars indicate SD (n = 10).

**Supplementary Figure S5.**
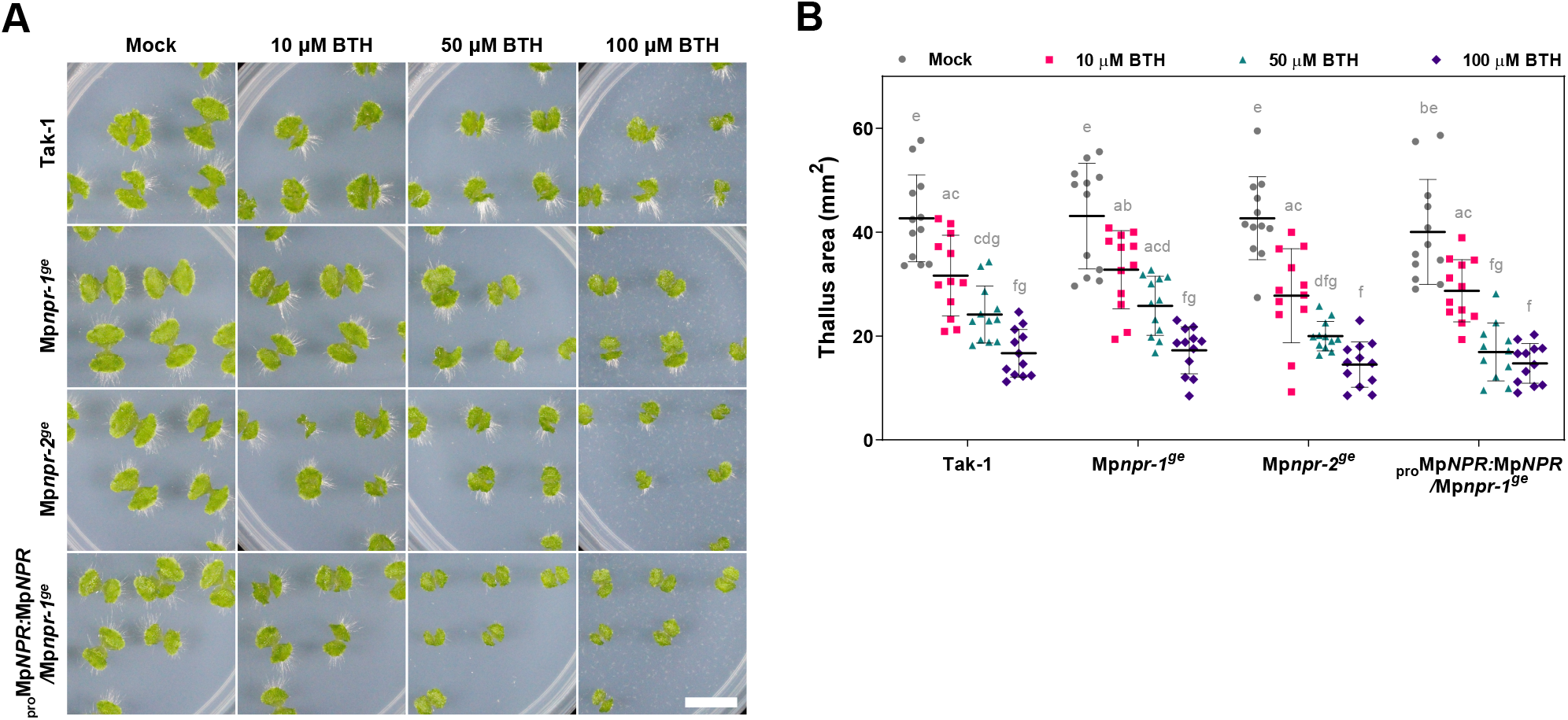
BTH sensitivity was unchanged in Mp*npr* mutants compared to Tak-1. (A) Ten-day-old *M. polymorpha* grown BTH conditions showing that BTH sensitivity was unchanged in Mp*npr* mutants compared to Tak-1. Scale bar, 1 cm. (B) Quantification of thallus projected area shown in A. Different letters represent statistical significances (Tukey’s test; *p* < 0.01). Error bars indicate SD (n = 12).

**Supplementary Figure S6.**
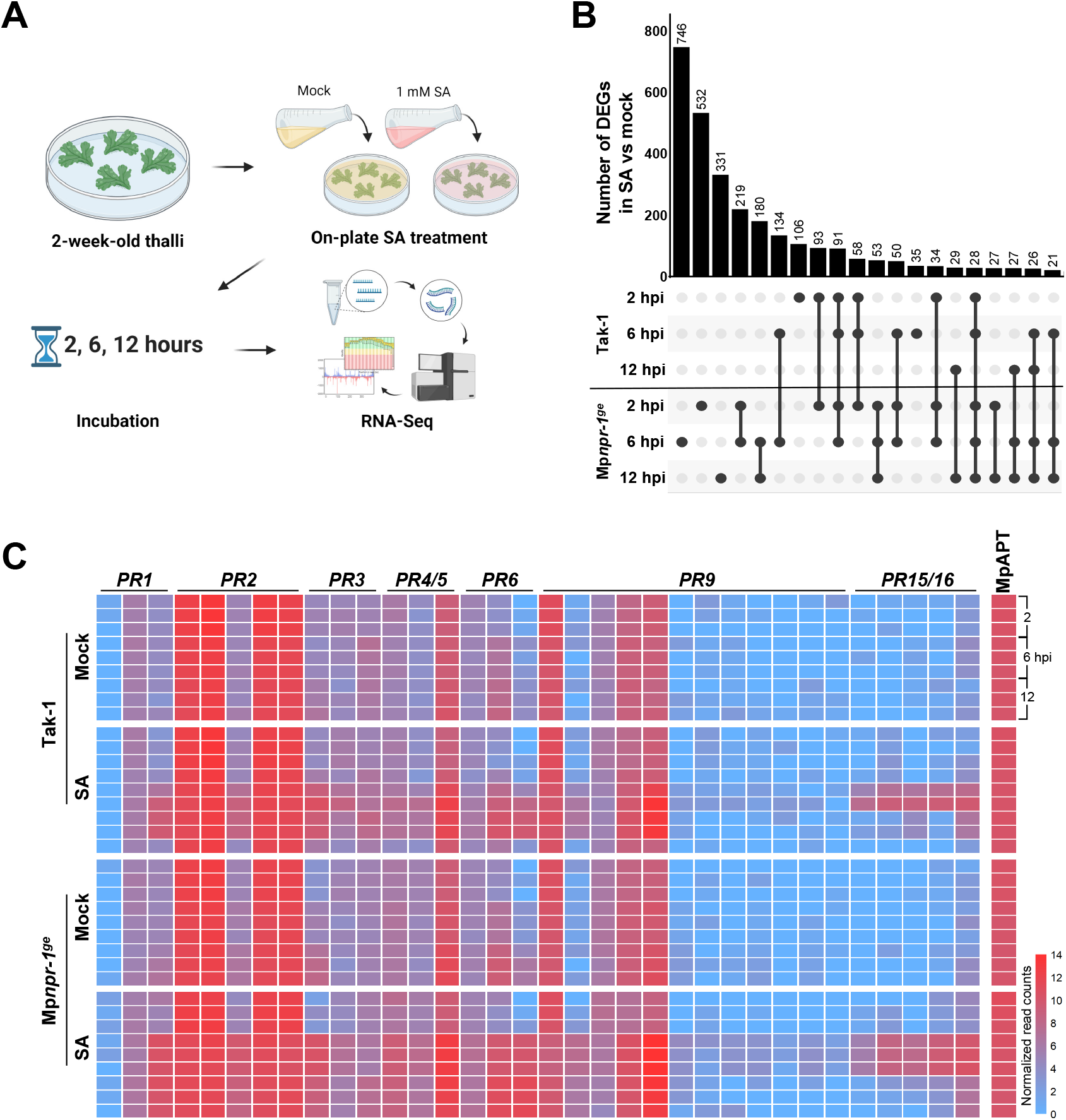
RNA-Seq analysis comparing Tak-1 and Mp*npr-1^ge^* thalli in Mock and SA conditions. (A) Schematic of thalli preparation for RNA-Seq analysis. (B) UpSet plot showing the number of unique and conserved DEGs (Mock vs SA, |Log2FC| ≥ 1, adjusted *p* < 0.05) in each dataset. Linked dots indicate that datasets share the DEGs. Comparisons showing more than 20 DEGs are shown. (C) Heatmap of selected Mp*PR* gene expressions in individual samples.

**Supplementary Figure S7.**
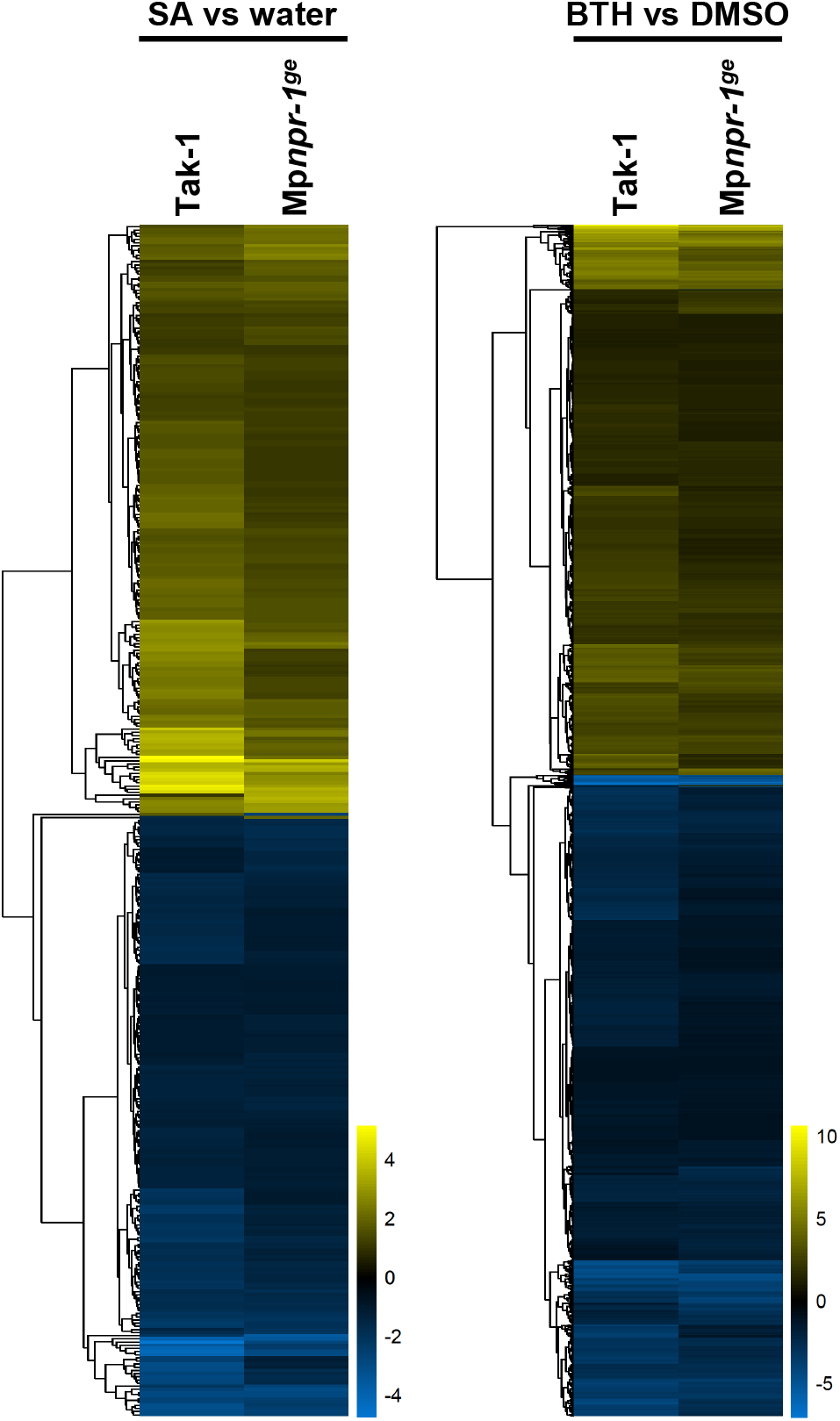
RNA-Seq analysis comparing Tak-1 and Mp*npr-1^ge^* gemmae in Mock, SA, and BTH conditions. Heatmap of DEGs (|log2FC| ≥ 1, adjusted p < 0.05) in Tak-1 and Mp*npr-1^ge^* gemmae upon SA (left) and BTH (right) treatments compared to each mock conditions.

